# Root phenolics as potential drivers of preformed defenses and reduced disease susceptibility in a paradigm bread wheat mixture

**DOI:** 10.64898/2026.01.13.699261

**Authors:** Laura Mathieu, Amandine Chloup, Soline Marty, Justin Savajols, Christine Paysant-Le Roux, Alexandra Launay-Avon, Marie-Laure Martin, Jean Chrisologue Totozafy, François Perreau, Adam Rochepeau, Claudia Rouveyrol, Pierre Pétriacq, Jean-Benoît Morel, Louis Valentin Méteignier, Elsa Ballini

**Affiliations:** PHIM Plant Health Institute, Univ Montpellier, INRAE, CIRAD, Institut Agro, IRD, Montpellier, France; Institute of Plant Sciences Paris-Saclay (IPS2), Université Paris-Saclay, CNRS, INRAE, Université Evry, Gif sur Yvette, France; Université de Paris, Institute of Plant Sciences Paris-Saclay (IPS2), 91190, Gif sur Yvette, France; Université Paris-Saclay, AgroParisTech, INRAE, UMR MIA Paris-Saclay, 91120, Palaiseau, France; Université Paris-Saclay, INRAE, AgroParisTech, Institut Jean-Pierre Bourgin for Plant Sciences (IJPB), Versailles, France; Bordeaux Metabolome, MetaboHUB, PHENOME-EMPHASIS, Villenave d’Ornon, France; Univ Bordeaux, INRAE, UMR1332 Biologie du Fruit et Pathologie, 33882 Villenave d’Ornon, France; PHIM Plant Health Institute, Univ Montpellier, CIRAD, Institut Agro, INRAE, IRD, Montpellier, France

**Keywords:** Metabolomics, multi-omics, plant-plant interactions, root exudates, Septoria tritici blotch, transcriptomics, wheat

## Abstract

Plant-plant interactions modulate foliar disease susceptibility in intraspecific mixtures. However, the molecular events including signals and responses underlying the reduction in disease susceptibility remain largely unexplored. Here, we developed an experimental system that can abolish root-mediated interactions between plants in a model of bread wheat varietal mixture. We then performed transcriptomic and metabolomic analyses to uncover the molecular responses linked to decreased susceptibility to Septoria tritici blotch in plant-plant interactions. Our analysis revealed that disrupting root chemical interactions impaired the reduction in susceptibility to Septoria and identified phenolic compounds as potential key mediators. The plant-plant interactions under study triggered significant molecular changes in specialized metabolism, biotic interactions, transporters, and responses to resources. Disrupting root interactions canceled both the macroscopic and molecular responses, thus providing a strong link between them. These insights provide a deeper understanding of the molecular basis of plant-plant interactions and the processes involved in reducing disease susceptibility in intraspecific mixtures.

**Significance statement:** Neighboring plants mediate resistance to leaf fungal pathogens by releasing root-derived molecules. These interactions trigger multi-omic reprogramming of defenses in both leaves and roots. Enhanced resistance in varietal mixtures is associated with the early activation of defense pathways.

Graphical abstract

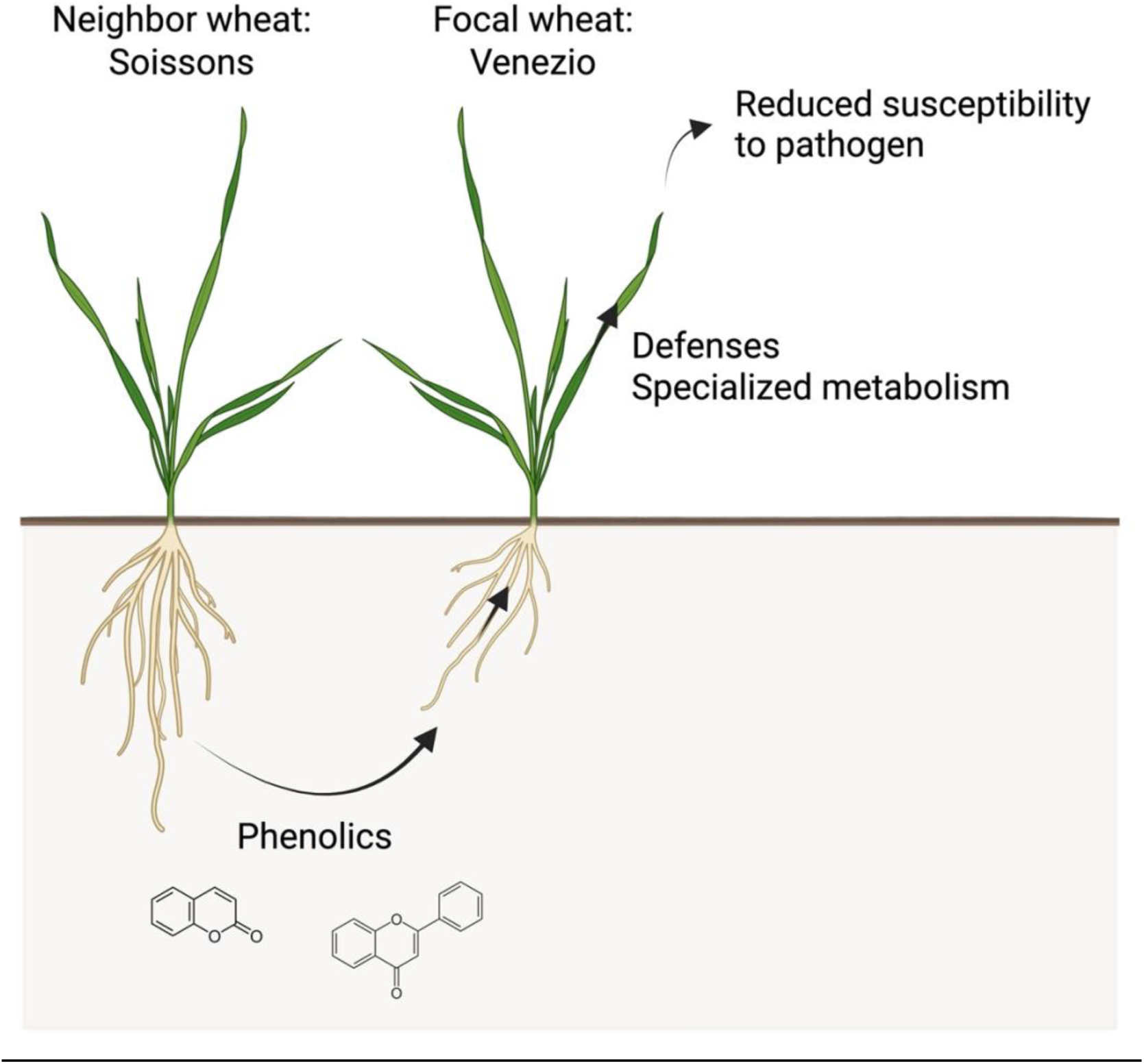

## Introduction

Plants continuously respond to their environment by modifying their physiology, which results in phenotypic changes. While the effects of biotic factors such as beneficial microbes and pathogens, as well as abiotic factors like light, temperature, and other microclimatic conditions, are well-documented (Mao *et al*., 2023), less attention has been given to the influence of neighboring plants. In natural and agricultural ecosystems, plants interact with each other through complex mechanisms, involving above- and belowground molecular signaling, plant-associated microbes, and resource availability (Subrahmaniam *et al*., 2018; Bilas *et al*., 2021; Mathieu, Ballini, *et al*., 2025). These plant-plant interactions can significantly influence a wide range of phenotypic traits, including above- and belowground development (Yang *et al*., 2015; Montazeaud *et al*., 2025) and susceptibility to diseases (Mathieu, Ballini, *et al*., 2025).

In addition to macroscopical responses induced by plant-plant interactions, more subtle and causative molecular changes also occur. However, the molecular mechanisms by which plants respond to neighboring individuals remain largely uncharacterized. To date, most research on plant-plant interactions has focused on interactions involving parasitic plants (Jhu and Sinha, 2022), weeds (Horvath *et al*., 2023), or intercropping systems where different species are mixed (Dai *et al*., 2018; Liu *et al*., 2021). Only a limited number of these studies have applied omic approaches in this context, with most focusing on transcriptomics in intercropping systems (Subrahmaniam *et al*., 2018; Horvath *et al*., 2023; Dai *et al*., 2018; Vazeux-Blumental *et al*., 2025). These studies have highlighted the involvement of oxidative stress signaling, defense responses, specialized metabolism (including flavonoids), light perception, photosynthesis, transport mechanisms, auxin signaling, and nitrogen-related pathways in responses to neighboring plants. Despite these diverse insights, integrative studies combining macroscopical and molecular analyses of plant-plant interactions remain rare (Subrahmaniam *et al*., 2018). In our previous work on enhanced susceptibility to *Zymoseptoria tritici* in a model mixture of durum wheat, we observed that the presence of conspecific neighbors altered the expression of tested immunity-related genes (Mathieu, Chloup, *et al*., 2025). Besides this report, the molecular mechanisms that drive phenotypic changes, especially those influencing reduced disease susceptibility in intraspecific mixtures, are still largely unexplored. Applying omic approaches to plant-plant interactions in intraspecific mixtures could unravel the molecular pathways underlying signaling between plants.

Intraspecific diversity in wheat varietal mixtures has been shown to generally reduce susceptibility to Septoria tritici blotch (STB), both in field (Kristoffersen *et al*., 2019; Montazeaud *et al*., 2022) and controlled conditions (Mathieu, Ducasse, *et al*., 2025). STB, caused by the fungal pathogen *Zymoseptoria tritici*, is one of the most devastating foliar diseases of wheat in Europe (Fones and Gurr, 2015). The plant response to STB is relatively well understood and involves basal immunity, salicylic acid and jasmonic acid signaling, and the biosynthesis of flavonoids, lignin, and other defense-related compounds (Rudd *et al*., 2015; Brennan *et al*., 2019; Seybold *et al*., 2020). Despite growing evidence that plant-plant interactions can reduce disease susceptibility, the molecular mechanisms by which they modulate susceptibility to STB remain unknown.

Recent studies highlight the importance of root-derived signals in mediating plant-plant interactions (Guerrieri and Rasmann, 2024). Several metabolites have been identified as signaling compounds in mono-genotypic or interspecific contexts, including strigolactones, loliolides, benzoxazinoids, flavonoids, and phenolic compounds (Mathieu, Ballini, *et al*., 2024). However, whether root-derived signals can modulate disease susceptibility in conspecific neighboring plants has not yet been demonstrated. Beyond the identification of specific signaling compounds, manipulating root-derived signals provides a valuable approach to functionally validate their role in modulating traits of interest. To this end, we developed a dedicated experimental setup that suppresses root chemical interactions, enabling us to assess the consequences of disrupted root signaling and to generate hypotheses explaining the associated modulation of disease susceptibility.

Here, focusing on a binary mixture of bread wheat (*Triticum aestivum*), we present a multi-omic analysis of plant-plant interactions that reduce susceptibility to STB. Using chemical-trapping barriers to alter root-root interactions between cultivars, we are able to more precisely link changes in STB susceptibility with underlying molecular responses. Through omic approaches, we aim to characterize the molecular responses of wheat plants to root exudates from neighboring individuals associated with reduced susceptibility.

## Results

### Reduced susceptibility to Septoria tritici blotch (STB) in wheat variety mixtures: The case of Venezio and Soissons

The susceptibility to Septoria tritici blotch (STB) was assessed across all possible binary mixtures of five bread wheat varieties. We observed 10/20 cases where STB susceptibility differed between mixed and pure stands (Figure S1). Notably, the variety Venezio displayed significantly reduced susceptibility to STB by ∼40% when grown in combination with Soissons, compared to its pure stand. This reduction was consistently observed across replicates (Figures 1 and S1), demonstrating the robustness and repeatability of this phenotype. As expected, RT-qPCR analysis confirmed that the reduced STB symptoms in the Venezio-Soissons mixture were associated with a decrease in the expression of pathogen housekeeping genes (Figure S2). To investigate the molecular and phenotypic determinants underlying reduced disease susceptibility in bread wheat mixtures, we concentrated our efforts on the Venezio-Soissons combination as a model system.

**Figure 1.**
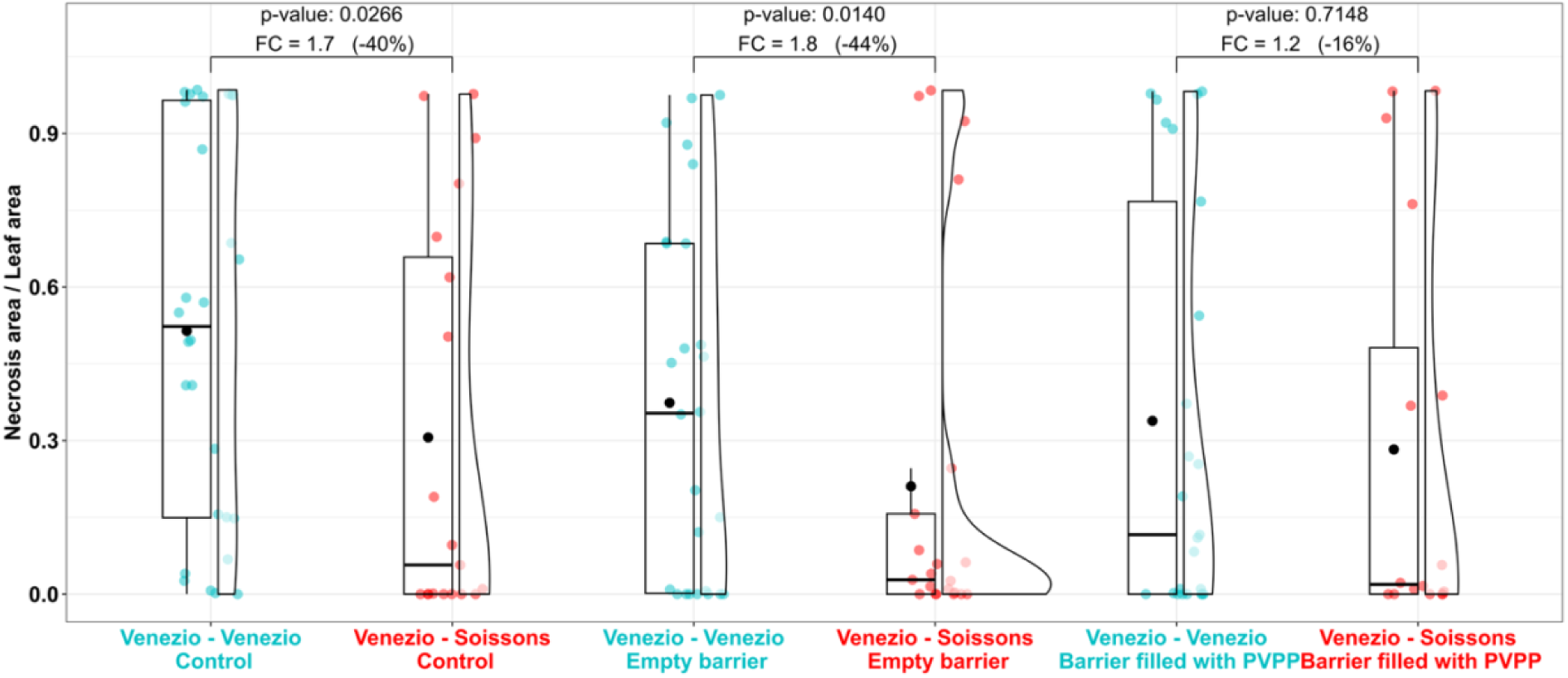
Effects of neighbor genotype and root barrier on Septoria tritici blotch symptoms in a bread wheat cultivar mixture. Septoria tritici blotch symptoms on the Venezio genotype were assessed by necrotic leaf area 17 days after inoculation. Venezio was grown in pure stands or mixed with Soissons, under control conditions or with either an empty barrier or a barrier filled with polyvinylpolypyrrolidone (PVPP) between genotypes. Statistical comparisons of Venezio symptoms between pure and mixed conditions were conducted using wilcoxon tests.

### Lower STB susceptibility associated with increased root intrusion in Venezio when grown with Soissons

To determine whether traits other than STB susceptibility are influenced by the interaction between Venezio and Soissons, we compared both aerial and root traits of Venezio in pure and mixed conditions. Our analysis indicated that Venezio displayed more intrusive roots when grown with Soissons compared to pure stands (Figure S3), suggesting that Venezio roots grow toward its neighbor Soissons. In contrast, no changes in aerial phenotypes were observed between the mixed and pure Venezio (Figure S3). These findings suggest that the Venezio-Soissons interaction not only impacts STB susceptibility but also root placement patterns.

### Phenolic root exudates could drive reduced STB susceptibility in the Venezio-Soissons mixture

To explore the molecules triggering the reduced STB susceptibility of Venezio when grown alongside Soissons, we hypothesized that molecular signals, specifically phenolics in root exudates, were involved in the interaction. To test this, we placed either a porous barrier between the two genotypes to hamper direct root contact or the same barrier filled with polyvinylpolypyrrolidone (PVPP), which can bind phenolic compounds (Folch-Cano *et al*., 2013; Durán-Lara *et al*., 2015). When the barrier was empty, STB symptoms were reduced in the mixture compared to the pure condition (Figure 1), as observed without barriers. However, the modulation of STB susceptibility was abolished when the barrier was filled with PVPP (Figure 1). Thus, belowground phenolic compounds likely mediate the observed reduction in STB susceptibility in Venezio when co-cultivated with Soissons.

### Coumarins and flavonoids: Candidate metabolites triggering a reduced STB susceptibility in the mixture

To identify the phenolic compounds effectively blocked by our experimental setup and potentially involved in the Venezio–Soissons interaction, we performed a metabolomic analysis of soil associated with Venezio grown in pure or mixed stands, using either empty or PVPP-filled barriers. Global metabolic profiles differed more strongly between mixed and pure conditions when the barrier was empty than when it was filled with PVPP (Figure 2a), and a significant neighbor × barrier interaction was detected on the first PLS-DA axis. Examination of individual metabolite patterns (Figure 2b) revealed a subset of compounds that accumulated differentially in mixtures relative to pure stands only in the presence of the empty barrier, but not when PVPP was included. The accumulation dynamics of these metabolites—particularly those in clusters S1 and S2 (Figure 2c)—were consistent with the phenotypic differences observed in STB symptom expression.

**Figure 2.**
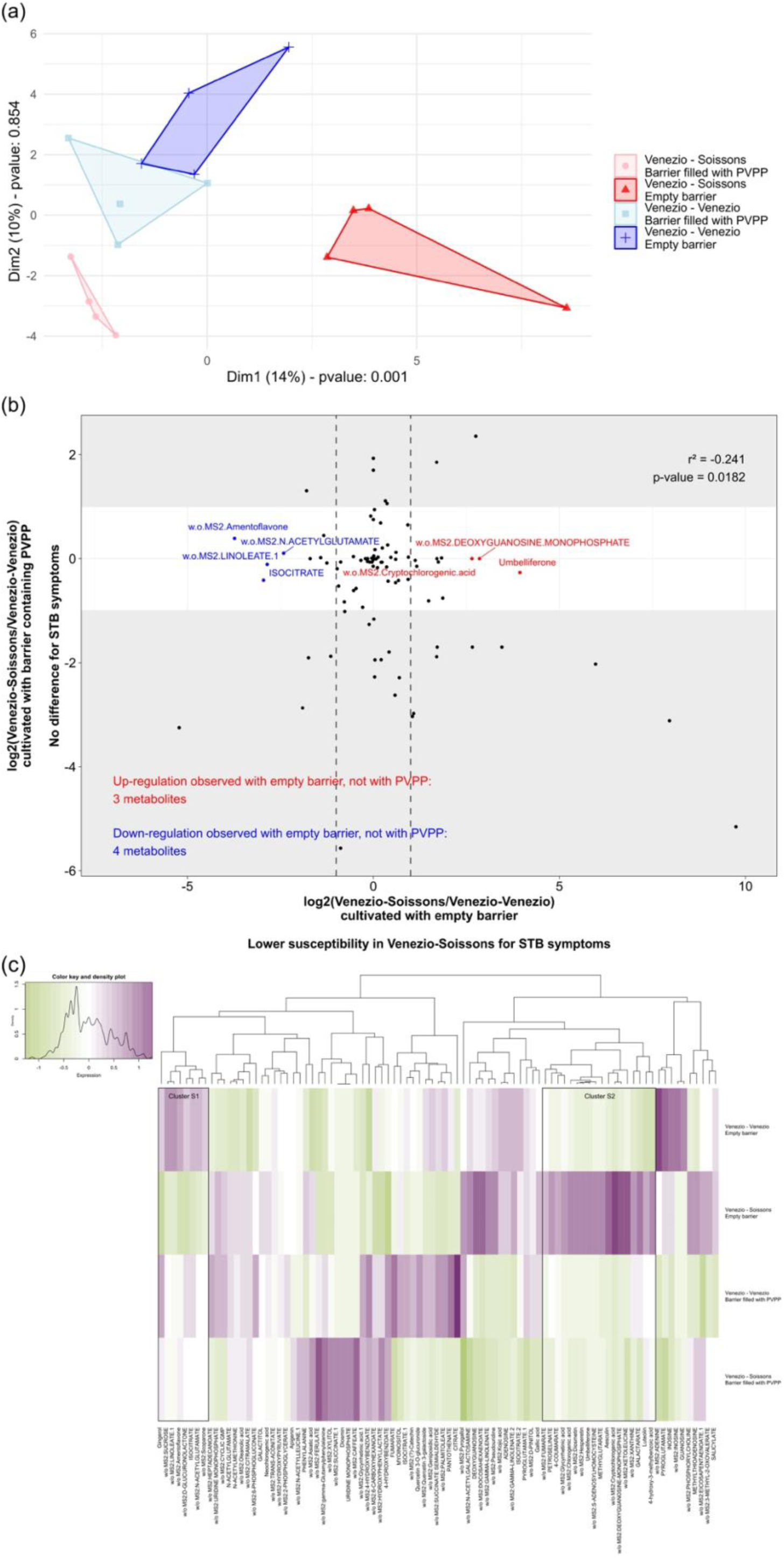
Effects of neighbor genotype and root barrier on soil metabolomic profiles in a bread wheat cultivar mixture. (a) Partial least squares discriminant analysis (PLS-DA) of metabolomic profiles from soil associated with Venezio grown in pure stands or mixed with Soissons with an empty barrier or a barrier containing polyvinylpolypyrrolidone (PVPP), at 21 days after germination (day of inoculation). Statistical analyses for each PLS-DA dimension were performed to test the interaction between the neighbor (Venezio *vs*. Soissons) and the barrier (empty *vs*. PVPP-filled barrier), using a non-parametric ANOVA. (b) Scatterplot and (c) heatmap of soil metabolites associated with Venezio grown in pure stands *versus* in a mixture with Soissons, under empty barrier or PVPP-filled barrier conditions. Metabolites were considered as significantly differentially accumulated if they exhibited a log2 fold-change less than −1 or greater than 1 and a z-score below −2 or above 2 in the empty barrier, while displaying a log2 fold-change between −1 and 1 and a z-score between −2 and 2 in the PVPP-filled barrier. The significantly up-accumulated metabolites with an empty barrier and not with PVPP were colored in red, while the down-accumulated ones were colored in blue.

To identify the phenolics effectively blocked by our experimental setup and potentially involved in Venezio-Soissons interaction, we conducted a metabolomic analysis of the soil associated with Venezio grown in pure and mixed stands, with either the empty or PVPP-filled barriers. A significant interaction between the neighbor and barrier effects was observed on the first PLS-DA axis, showing that the global metabolic profiles differed more strongly between mixed and pure condition with the barrier was empty, than when it was filled with PVPP (Figure 2a). Detailed accumulation profiles (Figure 2b) revealed metabolites that showed differential accumulation in the mixture, compared to the pure stand with the empty barrier, but not with a barrier filled with PVPP. The accumulation patterns of these metabolites (in particular clusters S1 and S2, Figure 2c) were consistent with the phenotypic differences observed in STB symptoms. Several phenolic compounds following this pattern were annotated, including coumarins and flavonoids (Table S1). One example is umbelliferone, a coumarin that showed significantly higher accumulation in the mixture with the empty barrier, and not in the PVPP-filled barrier condition (Figure 2b). These findings thus identify a limited set of metabolites amongst coumarins and flavonoids that were found in rhizochemicals and could explain reduced STB susceptibility.

### Mixture-induced molecular reprogramming in Venezio leaves and roots

We next investigated whether the alteration of root-mediated plant-plant interactions influences wheat molecular responses to its neighbor. For that matter, we performed transcriptomic and metabolomic analyses on leaves and roots of Venezio grown in pure stands or mixed with Soissons, under both empty and PVPP-filled barrier conditions, at 0 day post-inoculation (dpi). Our analysis of the molecular responses in the absence of infection was organized into two datasets – one for leaves and one for roots (Table S2), with PCA results confirming the reliability of our replicates through clear clustering of samples by condition (Figure 3). A distinct effect of neighbor (Venezio in dark blue *vs.* Soissons in dark red) was observed under the empty barrier condition across all datasets (Figure 3), indicating that both transcriptomes and metabolomes in the leaves and roots are reprogrammed in response to an intraspecific neighbor. In contrast, with PVPP-filled barriers, root transcriptomes and metabolomes no longer differed between pure and mixed conditions, while leaves still showed distinct profiles, but in a pattern different from that observed under the empty barrier condition (Figure 3). This demonstrates that disrupting root-mediated interactions alters molecular responses, consistent with phenotypic observations of STB symptoms, and that additional non-sequestered signals can still induce responses in Venezio leaves albeit not linked to the observed decrease of susceptibility. Altogether, these results confirm that the presence of a neighbor modifies transcriptomic and metabolomic profiles under the empty barrier condition, and that the alteration of root-root interactions is also evident at the molecular level.

**Figure 3.**
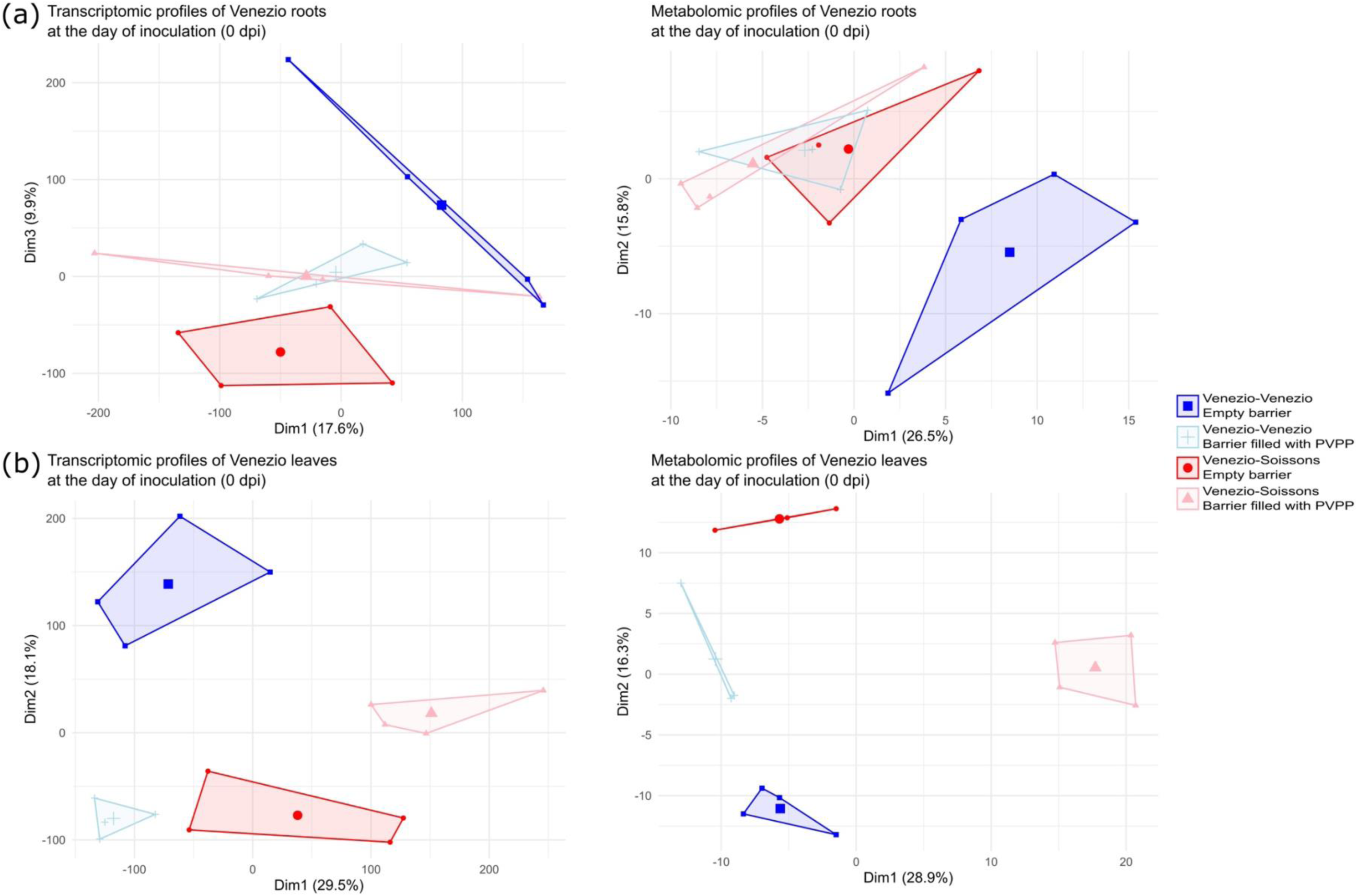
Transcriptomic and metabolomic profiles in pure or mixed Venezio roots and leaves when grown with an empty barrier or a PVPP-filled barrier. Principal component analysis (PCA) of transcriptomic (left panel) and metabolomic (right panel) profiles are shown for Venezio (a) roots and (b) non-inoculated leaves, grown in pure stands or mixed with Soissons, with an empty barrier or a barrier containing polyvinylpolypyrrolidone (PVPP), at 21 days after germination (day of inoculation).

### PVPP-filled barriers block most molecular changes observed between pure and mixed conditions with the empty barriers

In order to better qualify the molecular changes that were observed between mixed and pure conditions and altered by the PVPP-barrier, we next analyzed the differentially expressed genes (DEGs) and differentially accumulated metabolites (DAMs) between mixed and pure conditions under empty and PVPP-filled barrier conditions at 0 dpi. This analysis revealed that the majority of DEGs and DAMs identified in Venezio in response to an intraspecific neighbor under the empty barrier were not differentially regulated with the PVPP-filled barrier (Figure S4, Table S3), whether in roots or leaves.

First in roots, 1000 up-regulated genes in the mixed compared to pure Venezio were identified under the empty barrier condition, with 74% showing no differential expression with PVPP (Figure S4a), which indicates that the alteration of plant-plant interactions by PVPP cancels most neighbor-induced gene regulation in this tissue. Moreover, none of the 290 down-regulated genes showed differential expression with the PVPP-filled barrier (Figure S4a). Cluster profiling of DEGs under the empty barrier condition revealed distinct expression patterns, with clusters T1, T2 and T4 displaying altered gene expression only under the empty barrier (Figure S5a). Similarly, 94% of DAMs detected with the empty barrier were not regulated when PVPP was present (Figure S4a). This pattern was observed in clusters M2, M3, M4, M5 and M6 (Figure S6a).

Second in leaves, the differential regulation of 64% of DEGs and 48% of DAMs under the empty barrier condition was cancelled with PVPP (Figure S4b). Transcriptomic cluster analysis revealed that clusters T3, T5 and T6 displayed differences in gene expression under the empty barrier, which were not observed with the PVPP-filled barrier, while for metabolomic analysis, all clusters except cluster M6 followed this same pattern (Figure S6b).

Overall, the global cancellation of neighbor-induced molecular changes by PVPP was consistent across both tissues and echoes the loss of STB phenotype.

### Potential molecular signatures responsible for the phenotypes triggered by intraspecific neighborhood

To further identify the molecular changes observed in the presence of an intraspecific neighbor that depend on phenolics, we then identified DEGs or DAMs between the mixture and pure stand, present in the empty barrier condition and absent in the PVPP-filled barrier condition at 0 dpi (Figure S7, Table S3).

This analysis revealed 586 increased and 272 decreased transcripts in the mixture for roots, and 616 increased and 1097 decreased transcripts for leaves (Figure 4 – left panels, Tables S3-5). In roots, up-regulated genes were enriched in pathways related to metabolism of terpenes and carotenes, biosynthesis of oxylipins and phenylpropanoids, nitrogen responses, and biotic interaction processes. These included genes encoding carotenoid cleavage dioxygenase-like proteins, flavanone dioxygenases, peroxidases, glutathione S-transferases, and receptor-like protein kinases. In contrast, down-regulated genes were enriched for regulation of root development, responses to wounding, catabolic functions and transporter activity (Figure 4a – right panel). In leaves, up-regulated genes – such as those encoding receptor-like protein kinases, WRKY transcription factors, Mildew Resistance Locus “O” (MLO)-like proteins and phosphatases – were enriched for functions associated with phosphorylation and responses to micro-organisms. Down-regulated genes, on the other hand, were enriched in pathways involved in light responses, transporters, nitrogen responses, chemical compound detoxification, and heme-containing molecule biosynthesis (Figure 4b – right panel). Notably, this group included several cytochrome P450 genes.

**Figure 4.**
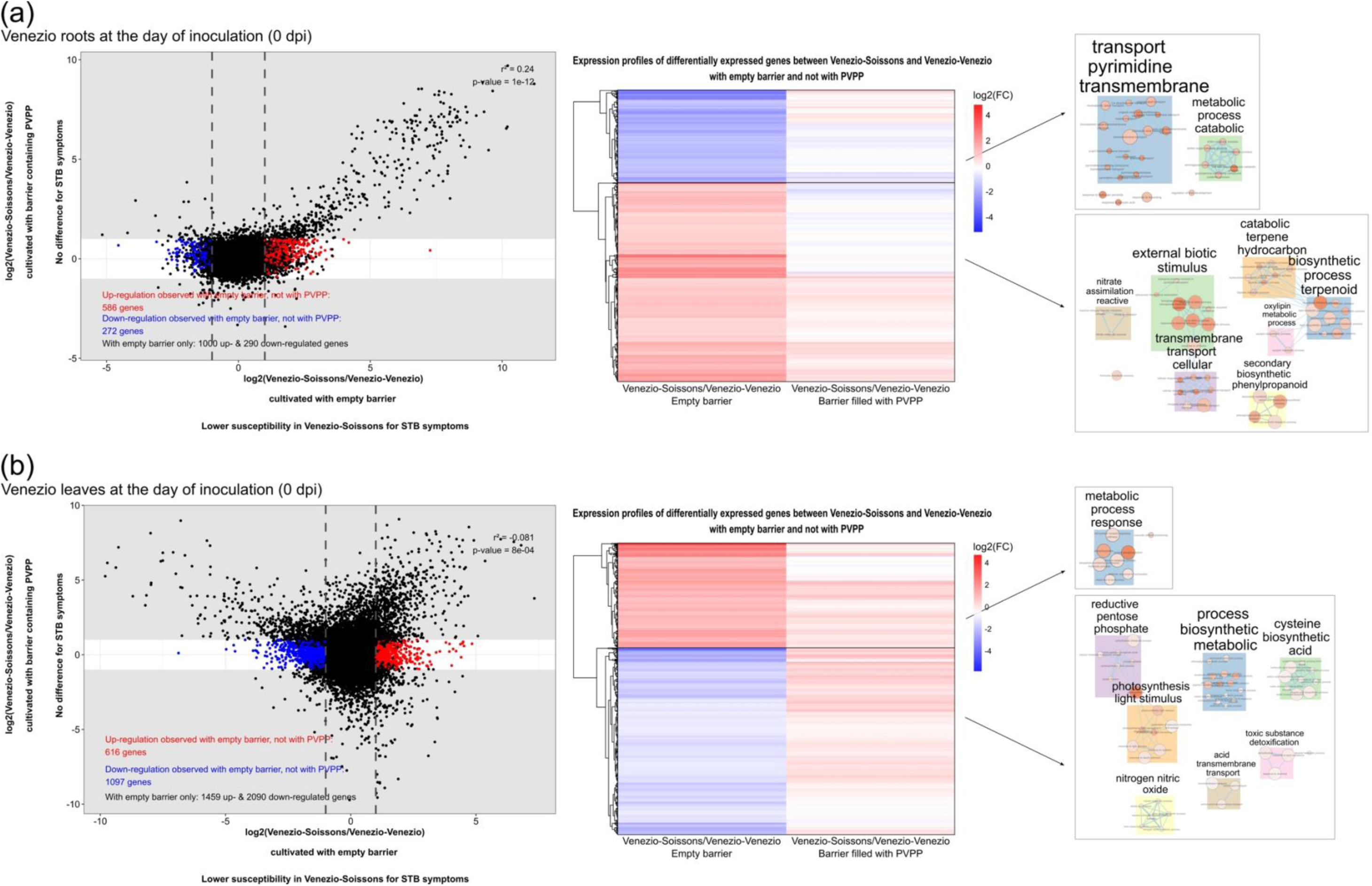
Expression profiles and Gene Ontology (GO) enrichment results of differentially expressed genes between pure or mixed Venezio when grown with an empty barrier, and not with a PVPP-filled barrier. Expression profiles (left panels) and enriched biological processes (right panel) are shown for differentially expressed genes between pure and mixed conditions with Soissons, when grown with an empty barrier, and not with a barrier containing polyvinylpolypyrrolidone (PVPP), in Venezio (a) roots and (b) leaves at 21 days after germination (day of inoculation). In the right panel, circle color indicates significance (red = more significant) and circle size represents the number of genes associated with each GO term. Genes were considered significantly differentially expressed if they showed a log2 fold-change of less than −1 or greater than 1 and an adjusted p-value lower than 0.05 in the empty barrier condition, while exhibiting a log2 fold-change between −1 and 1 and an adjusted p-value higher than 0.05 with the PVPP-filled barrier. GO term enrichment analysis was performed using a hypergeometric test followed by Bonferroni correction.

For metabolites, we identified 9 increased and 5 decreased metabolites in roots, and 8 increased and 5 decreased metabolites in leaves (Figure S8 – left panel, Tables S3 & S6-7). In roots, up-regulated metabolites included a benzene, a phenol ether, three lignan glycosides and two carboxylic acids, while down-regulated ones included a cinnamic acid and two carboxylic acids (Figure S8a – right panel). In leaves, up-regulated metabolites were two flavonoids and two carboxylic acids, with down-regulated ones including a cinnamic acid (Figure S8b – right panel).

These transcriptomic and metabolomic changes reveal molecular signatures of root-derived, molecule-mediated plant-plant interactions, in pathways potentially contributing to the observed differences in STB symptoms.

## Discussion

Thanks to an experimental design that prevents plant-plant interactions at the root level, this study has identified potential cues and responses that could contribute to decreasing the susceptibility of wheat to Septoria tritici blotch in mixtures (Figure 5). Indeed in the Venezio-Soissons interaction, phenolic compounds, particularly coumarins and flavonoids, appear to act as potential roots signaling molecules mediating the reduced STB susceptibility in the Venezio plants. We also identified molecular changes, before inoculation, in processes related to specialized metabolism, light and nitrogen responses, transporters, and biotic interactions that may be crucial in driving the observed symptom differences in the mixture.

**Figure 5.**
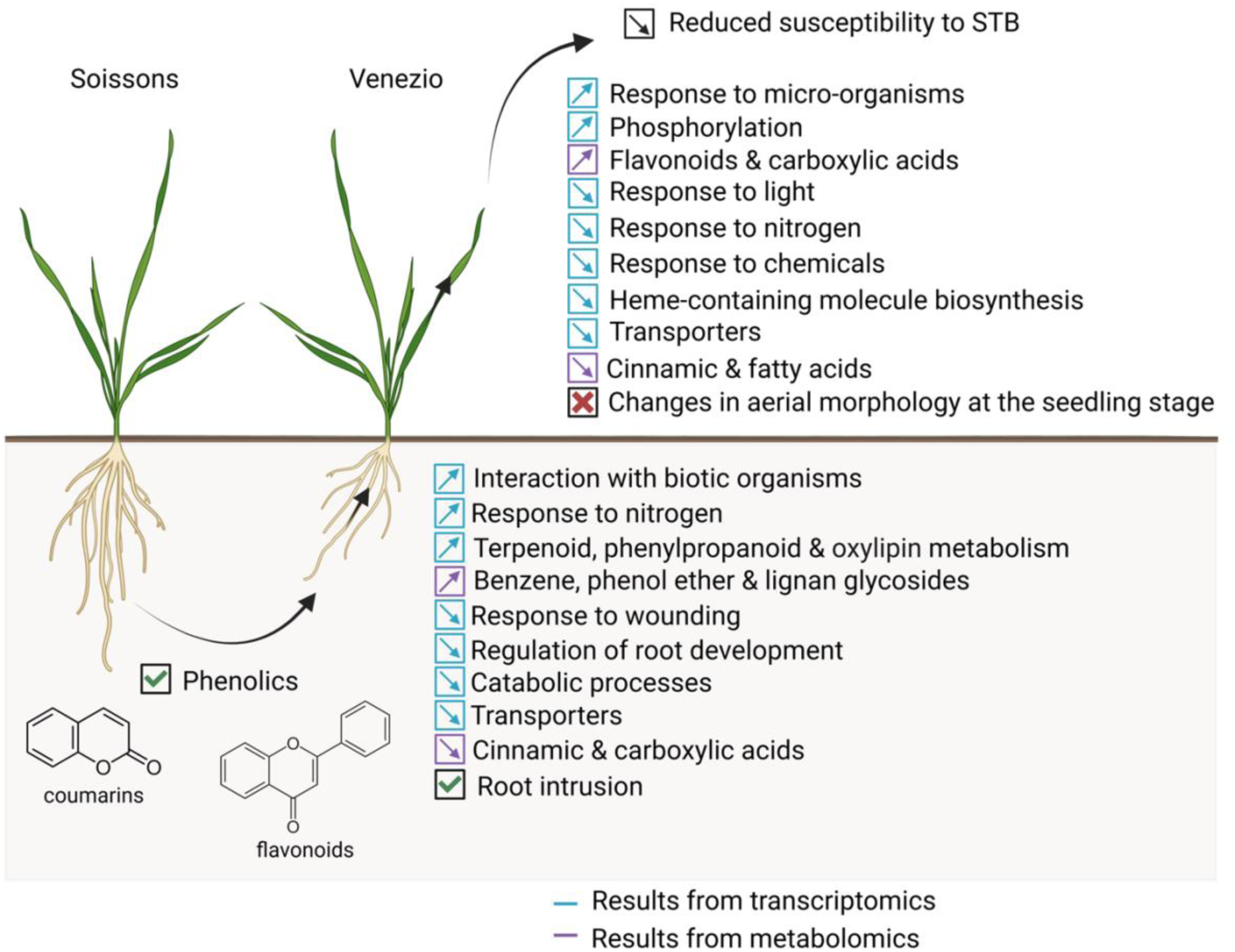
Scheme of the proposed signaling cascade in a bread wheat varietal mixture. In the Venezio-Soissons mixture, phenolics—specifically coumarins and flavonoids—trigger changes in gene expression and metabolite accumulation. In roots, these molecular shifts were associated with up-regulation of specialized metabolism pathways, nitrogen responses and biotic interaction processes, and down-regulation of root development, transporter activity or primary metabolism. In leaves, modifications include up-regulation of phosphorylation, flavonoid accumulation or response to oomycetes, and down-regulation of light responses, transporter genes, heme-containing molecule biosynthesis or nitrogen response. These molecular changes likely contribute to the reduced susceptibility to STB observed in this varietal mixture.

Phenolics, particularly flavonoids, and benzoic acids are known soil signaling molecules involved in plant-plant interactions (Mathieu, Ballini, *et al*., 2024). Among them, phenolics are particularly promising candidates for modulating disease susceptibility due to their well-documented role in plant defense (Kumar *et al*., 2020) and their involvement in interspecific interactions (Xie *et al*., 2023). In this study, our analysis of soil metabolome suggests that such compounds may also contribute to the modulation of STB susceptibility within intraspecific mixtures. Notably, coumarin, one of our candidate phenolics, has been reported to induce oxidative stress by increasing H_2_O_2_ content and to down-regulate genes related to light responses (Araniti *et al*., 2017), while also enhancing resistance to pathogens through SA-signaling pathways (Vismans *et al*., 2022). These reported effects parallel our observations in the Venezio plants cultivated with Soissons, where we detected a down-regulation of light-responsive genes, and an induction of specialized metabolism and biotic stress-responsive genes. Such responses align with the known role of reactive oxygen species, including H_2_O_2_, in mediating plant responses to biotic stresses (Tyagi *et al*., 2022), as well as with the involvement of specialized metabolic pathways in defense activation (Delplace *et al*., 2020). Consistent with reduced susceptibility of Venezio in the presence of Soissons, we detected elevated leaf defense gene expression at the time when infection is performed (see below for details).

While phenolics have so far been associated with negative allelopathic effects in interspecific interactions (Xie *et al*., 2023), this study provides an example that they also underlie positive allelobiotic effects in intraspecific plant-plant interactions. Collectively, these results support the hypothesis that neighboring plants influence molecular responses in Venezio *via* phenolic compounds, thereby enhancing its defense mechanisms and reducing susceptibility to pathogens.

It is also plausible that some of these phenolics, or their derivatives, not only act externally in the rhizosphere but may also function as endogenous signals within the plant itself. Indeed, the metabolites detected in the Venezio roots, which are directly or indirectly linked to phenylpropanoid pathways and phenolic compounds (Fraser and Chapple, 2011), raise the possibility that one of these molecules may play a signaling role internally. However, further validation is required to confirm this hypothesis. The perception and transduction of signaling molecules in plant-plant interactions remains one of the least understood areas in the field, and more research is needed to elucidate these mechanisms.

Although we did not directly test the role of phenolics in the observed root phenotype of intrusion, it is noteworthy that previous work has shown that allelochemicals can alter root placement in mixed plantings, notably in wheat (Wang *et al*., 2023). This suggests that the phenolics identified here may shape root responses in our system.

This significant increase in root intrusion observed in the mixed Venezio plants, in response to potential cues, was accompanied by the down-regulation of genes associated with root development, pointing to a shift in developmental regulation potentially linked to altered resource foraging or neighbor perception (Fang *et al*., 2013).

These results suggest that root exudates may function not only as signaling molecules that trigger aboveground responses but also as regulators of belowground developmental plasticity in intraspecific interactions.

The reduced disease susceptibility observed in intraspecific mixtures appears to reflect a broader reprogramming of molecular pathways related to plant defense. Under the empty barrier condition only, we observed an up-expression of genes typically involved in responses to micro-organisms in the leaves of Venezio grown with an intraspecific neighbor, as well as genes related to biotic interactions in roots. Systemic acquired resistance, a central component of plant biotic stress response, has previously been reported to be induced under mixed plantings in response to secondary metabolite signals (Riedlmeier *et al*., 2017; Subrahmaniam *et al*., 2018). However, previous work testing a limited set of genes in an intraspecific mixture did not detect such enhanced immunity (Pélissier *et al*., 2021). Our findings therefore demonstrate that immunity can be constitutively modified in wheat mixtures, supporting the idea that the presence of the Soissons genotype induces a generalized biotic stress alertness in Venezio, potentially mimicking microbe recognition and thereby enhancing defense responses.

In parallel to immunity, genes related to phosphorylation processes were also up-regulated in the Venezio plants grown with Soissons. Protein phosphorylation is a central mechanism of plant signaling cascades, particularly under stress conditions (Zhang *et al*., 2023). It is essential for activating immune responses, which mediate the perception of external cues or activate downstream responses (Erickson *et al*., 2022). These signaling pathways play a central role in damage-triggered immunity (Tanaka and Heil, 2021) and are implicated in mediating resistance against STB in wheat (Brennan *et al*., 2019).

Our transcriptomic and metabolomic analyses also revealed a significant enhancement of specialized metabolic pathways in the mixture, particularly those related to flavonoids in leaves and terpenoids, oxylipins, phenylpropanoids, benzenoids, and lignan glycosides in roots. These compounds are well-established contributors to plant defense (Delplace *et al*., 2020). Flavonoids are well-characterized defense compounds that contribute to pathogen resistance, including resistance to STB (Seybold *et al*., 2020; Ramaroson *et al*., 2022). Together, these molecular patterns indicate that intraspecific plant-plant interactions reprogram by enhancing constitutive expression of defense-related networks, offering a powerful mechanism for natural disease suppression.

Altogether, our study sheds new light on the molecular basis of intraspecific plant-plant interactions and identifies root-exuded phenolic compounds as key allelobiotic signals that can modulate disease susceptibility in leaves. By demonstrating that plants can constitutively induce defenses in response to chemical cues from intraspecific neighboring plants, our work contributes to a better understanding of how plants interact with each other. These findings also open promising avenues for breeding or engineering plants to develop sustainable agricultural systems that harness beneficial plant-plant interactions, improving resilience and reducing reliance on chemical inputs.

## Materials and Methods

### Plant material

The selected bread wheat genotypes analyzed in this study are varieties listed in the French Catalogue and were chosen for their susceptibility to *Zymoseptoria tritici*. This criterion enabled us to study changes in basal immunity rather than resistance genes, as in (Pélissier *et al*., 2023). A total of 5 bread wheat varieties were included in the analysis.

### Plant growth conditions

The wheat plants were cultivated in a greenhouse at PHIM (Montpellier, France) under controlled conditions. The photoperiod was 16 hours of light and 8 hours of darkness, with a temperature of 24°C/20°C and a light intensity of 250 μmol/s/m2.

Plants were grown in 8cm × 8cm × 8cm pots filled with a soil mixture consisting of 50% topsoil, 50% Neuhaus N2 soil, and 133.3 g of TopPhos for 100L soil. Each pot contained two rows: one with four plants of the focal genotype, on which phenotype is observed, and the other with four plants of the neighboring genotype. Bread wheat plants, either grown in pure stands or in mixtures, were cultivated under control conditions or with either an empty porous barrier or the same porous barrier containing polyvinylpolypyrrolidone (PVPP) to block the exchange of chemical molecules, particularly phenolics (Folch-Cano *et al*., 2013; Durán-Lara *et al*., 2015), between genotypes (Figure S9).

For root phenotyping, plants were cultivated in rhizoboxes (20 cm × 40 cm × 2 cm) filled with the same soil mixture, enabling detailed observation of root architecture. The rhizoboxes were inclined at a 30° angle relative to the horizontal plane to encourage root development along the lower face of the box. The sides of the rhizoboxes were shielded from light to prevent unwanted light effects on root development. In each rhizobox, two plants of the same genotype (pure) or two plants of different genotypes (mixture) were sown against the glass panel, with a spacing of 5 cm between them.

### Inoculation and symptom assessment

The focal genotypes were inoculated with the *Zymoseptoria tritici* strain IPO9415 by applying an inoculum of 10⁶ spores/mL in water containing Tween 20 to the last ligulated leaf using a paintbrush, 21 days after germination. Following inoculation, plants were placed in transparent plastic bags for three days to maintain high humidity and promote pathogen growth. Seventeen days post-inoculation, symptoms of Septoria tritici blotch (STB) were assessed using an Epson Perfection V370 Photo scanner. The SeptoSympto tool (Mathieu, Reder, *et al*., 2024) was used to quantify the surface area of necrosis, which refers to the lesions.

### Root and aerial trait assessment

Root and aerial traits were evaluated at different growth stages under both pure and mixed conditions (Figure S9). Root system architecture was imaged 21 days after germination, allowing clear differentiation between genotypes and enabling precise measurement of root angles for the Venezio genotype. Root growth angle (RGA) was quantified using ImageJ software by measuring the angle formed between the two outermost seminal roots. In addition to RGA, we measured the internal angle (IA), external angle (EA), and intrusion value, based on a vertical reference line extending 7 cm below the hypocotyl. The internal angle was defined as the half-angle on the side facing the neighboring plant, while the external angle was taken on the opposite side. The intrusion value was calculated as IA − EA; negative values indicate an escape phenotype (root growth away from the neighbor), whereas positive values reflect an intrusive phenotype (root growth toward the neighbor).

Aerial traits of the Venezio genotype were measured 38 days after germination. Plant height, leaf number, and tiller number were scored. Chlorophyll content was also assessed on the adaxial surface of the last ligulated leaf using a Dualex sensor (Goulas *et al*., 2004), providing insights into photosynthetic potential and plant health.

### Sample collection

Leaf samples were collected from the genotype Venezio, with each sample comprising a pool of the last expanded leaves from three different pots (12 leaves total per sample). Root samples were taken from the same three pots. After grinding, samples were divided for transcriptional and metabolomic analyses. Root and leaf samples were collected at 21 days after germination (day of inoculation) from uninoculated Venezio plants grown in pure stands or mixed with Soissons, with an empty porous barrier or a porous barrier containing PVPP between genotypes. For leaves, the central sections were cut into 0.5 cm pieces, while for roots, the tips were cut into 0.5 cm sections. The samples were immediately stored in liquid nitrogen.

Soil samples were taken at 21 days after germination (day of inoculation) from soil associated with Venezio in pure stands or mixed with Soissons (Figure S9), with an empty porous barrier or a PVPP-filled barrier separating the genotypes. Each soil sample consisted of a pool of three subsamples collected from the left, right, and center of the pot, along the line of Venezio plants.

### Soil metabolomic analysis

Metabolomics was performed from robotized sample preparations (Dussarrat *et al*., 2022). Briefly, five mg of dried soil metabolite extracts extracted in 2 x 300 µL (ethanol/water 80% v/v, Formic acid 0.1% v/v, Methyl vanillate 250µg/mL), sonicated (15 min, 4°C), centrifuged (5 min, 10 600 g, 4°C) and filtered (sterile 0.22µm Durapore membrane, Merck), were then analyzed by UHPLC-Q-Exactive Orbitrap MS/MS-Based Untargeted Metabolomics at *Bordeaux Metabolome* as described previously (Dussarrat *et al*., 2025). The LCMS sequence of 26 injections was randomized and comprised 18 biological samples (n = 3; 5 genotypes + 1 bulk soil control), 2 extraction blanks (no biological material), and 6 QC samples prepared by pooling 50 µL from each sample and standard. QCs corrected signal drift and enabled calculation of feature CVs to retain only robust signals. Data were processed via DDA MS2 deconvolution in MS-DIAL v4.9 with optimized parameters (Tsugawa *et al*., 2015). Annotations were performed based on MS1 spectra and MS2 DDA fragmentation information using in-house and FragHub HRMS/MS databases, including thousands of phytochemicals (Dablanc *et al*., 2024). Thus, putative annotation of differentially expressed metabolites resulted from MS-DIAL screening of the MS1 detected exact HR m/z and MS2 fragmentation patterns (Tsugawa *et al*., 2015). Additionally, the InChiKeys of annotated features were employed within ClassyFire to generate a structural ontology for chemical entities (Djoumbou Feunang *et al*., 2016). From 6369 raw metabolomic signals, 1168 LC–MS features were retained after curation (SN > 10; CV QC < 30%) and standardised to dry extract mass. Of these, 1104 were unmatched (no MS1/MS2 ID), 32 were putative metabolites (MS1 match only, level 3), and 64 were annotated metabolites (MS1 + MS2 level 2). All features were Pareto-scaled for normalization and analyzed via univariate and multivariate methods (PCA, PLS-DA, clustering) in MetaboAnalyst v6 (Pang *et al*., 2021).

### RNA extraction

Total RNA was extracted using the protocol described in (Delteil *et al*., 2012). Initially, frozen leaf tissues were ground in liquid nitrogen. Cells were lysed by adding 1 mL of Tri-reagent to the powdered sample. Nucleic acids were then separated by adding 200 µL of chloroform, incubating for 10 minutes at room temperature, followed by centrifugation at 13000g for 15 minutes at 4°C. The upper aqueous phase was collected, and 560 µL of isopropanol was added to precipitate the nucleic acids overnight at −80°C, followed by centrifugation at 13000g for 20 minutes. The resulting pellet was washed twice with 500 µL of 70% ethanol and centrifuged at 13000g for 5 minutes at 4°C, and then air-dried for 10 minutes before resuspension in 50 µL of ultrapure water on ice for 1 hour. DNase treatment was performed on 50 µL of RNA using the TermoFisher kit, followed by purification. 200 µL of RNA was mixed with 200 µL of phenol:chloroform:alcohol (25:24:1), and centrifuged at 16000g for 10 minutes at 4°C. The upper phase was collected and precipitated with 18 µL of 3M sodium acetate and 495 µL of pure ethanol at −80°C for 1 hour, followed by centrifugation at 16,000g for 30 minutes. The pellet was washed with 1 mL of 70% ethanol, centrifuged for 5 minutes, dried, and resuspended in 50 µL of ultrapure water. RNA quality and quantity were assessed using a Nanodrop spectrophotometer and verified for integrity via 2% agarose gel electrophoresis. The absence of DNA contamination was confirmed by qPCR using the Promega kit with EGM34 and EGM35 primers. The RNA were finally diluted to 0.2 µg/µL.

### Primers for the Zymoseptoria tritici RT-qPCR

Gene expression of two *Zymoseptoria tritici* housekeeping genes was analyzed in inoculated Venezio leaves, either grown in pure stands or mixed with Soissons, at 7 days post-inoculation (biotrophic phase) and 14 days post-inoculation (necrotrophic phase) (Figure S9). The primers used for RT-qPCR are listed in Table S8.

### Leaf & root transcriptomic analysis

#### Library construction

First, total RNA was purified using the RNA Clean & Concentrator Kits (Zymo Research). The integrity of the extracted RNA was controlled on Agilent RNA Nano chip, ref #5067-1511 (Agilent) and their quantification was performed with Quant-iT™ RiboGreen, ref #R-11491 (Thermo Fisher Scientific Inc). Second, RNA-seq libraries were constructed with 250 ng of total RNA using the Illumina Stranded mRNA Prep Ligation kit ref #20040534 (Illumina) at half volumes. Libraries were checked on Agilent DNA HS chip, ref #5067-4627 (Agilent) and quantified with Quant-iT™ PicoGreen™ dsDNA Reagent, ref #P7581 (Thermo Fisher Scientific Inc) to generate an equimolar multiplexed pool of libraries. The libraries were sequenced by the Genoscope in paired-end (PE) mode with 150 bases for each read on a NovaSeq6000 (Illumina) to generate between 15 and 37 millions of PE reads per sample.

#### Data preprocessing and mapping

First, to remove the least reliable data from the analysis, the raw data were filtered to remove any clusters that have “too much” intensity corresponding to bases other than the called base. The purity of the signal from each cluster was examined over the first 25 cycles and calculated as Chastity = Highest_Intensity / (Highest_Intensity + Next_Highest_Intensity) for each cycle. The default filtering implemented at the base calling stage allows at most one cycle that is less than the Chastity threshold (0,6). Second, adapter and primer sequences were removed on the whole read, along with nucleotides at both ends with a quality score lower than 20. Third, any sequences between the second unknown nucleotide (N) and the end of the read were discarded and reads shorter than 30 nucleotides were excluded from the dataset. Finally read pairs that come from the low-concentration spike-in library of Illumina PhiX Control were removed, and ribosomal RNA sequences were also eliminated using sortMeRNA (v2.1). To improve the final mapping pourcentage, reads were adapter trimmed again using BBduk from the BBmap suite (v38.84) with the options k=23 ktrim=r qtrim=r useshortkmers=t mink=11 trimq=20 hdist=1 tpe tbo minlen=30 using adapter sequences file included in the BBTools package.

Reads were mapped and counted using STAR v2.7.3 (Dobin *et al*., 2013) with the following parameters --alignIntronMin 5 --alignIntronMax 60000 --outSAMprimaryFlag AllBestScore --outFilterMultimapScoreRange 0 --outFilterMultimapNmax 20 on the *Triticum aestivum* reference genome (v2.13(A/B/D)x7) and its associated GTF annotation file (v2.1). Between 71.2 and 76.8 % of the reads were associated to annotated genes.

### Leaf & root metabolomic analysis

#### Extraction of polar and semi-polar specialized metabolites

Polar and semi-polar metabolites were extracted as described below. Samples consisted of 5 mg of dried and ground tissues. Briefly, 1 mL of methanol/water(50/50) + 0.1% of formic acid were added to each sample. The mixtures were sonicated (Fischer Scientific FB15050) for 15 min, then shaken in 2-mL tubes using a ThermoMixer™ C (Eppendorf) at 1400 rpm for 60 min and 25°C. The tubes were centrifuged at 14000 rpm for 10 min at 25°C; the supernatants were collected in glass tubes. Next, 1 mL of extraction solution was added to the 2mL tube containing the pellet, and the extraction procedure was repeated. The two supernatants were pooled and dried using a vacuum concentrator (SpeedVac). The dried extracts were resuspended in 100 µL acetonitrile:water (1:9) of ULC/MS grade. The resuspended extracts were filtered with a glass microfiber filters (Cat. NO. 1820-037, Whatman International Ltd., UK) and distributed in HPLC vials. A Quality Control (QC) made up of 5 µL from each sample by type of tissue was prepared in a separate HPLC vials.

#### UPLC-MS/MS analysis of specialized metabolites and UPLC-MS/MS data processing

Untargeted metabolomic analysis was performed using an Ultimate 3000 UHPLC system (Thermo Fisher Scientific) coupled with a QTOF Impact II quadrupole time-of-flight mass spectrometer (Bruker Daltonics, Bremen, Germany). The liquid chromatography and mass spectrometry conditions are described in (Boutet *et al*., 2022), except for a modification to the electrospray source temperature, which was set to 250 °C.

The UPLC-MS/MS data was processed using the pipeline described in (Boutet *et al*., 2022) with modifications. Briefly, .d data files (Bruker Daltonics, Bremen, Germany) were converted to .mzXML format using the MSConvert software (ProteoWizard package 3.0). A batch script containing all the steps—mass detection, chromatogram builder, deconvolution, deisotoping, alignment, and blank subtraction—was created in MZmine 4 (https://github.com/mzmine/mzmine/releases) using the parameters described in (Boutet et al., 2022). The script was then run on positive and negative data separately. Metabolite accumulation was normalized according to the weight of tissue used for the extraction. Pooled QC sample injections across LC-MS/MS run were used to evaluate the quality of the run and untargeted metabolomic dataset. The relative standard deviation across the QC samples was calculated for each metabolic feature detected. The metabolic features were filtered according to their QC variation and only metabolic features with QC variation < 25% were kept.

#### Molecular network and annotation of untargeted metabolomic data

As described previously, molecular networks were generated with MetGem 1.4.1 software (https://metgem.github.io) using the .mgf and .csv files obtained with MZmine 4 analysis. The molecular network was optimized for the ESI+ and ESI-datasets, and different cosine similarity score thresholds were tested. ESI- and ESI+ molecular networks were both generated using a cosine score threshold of 0.75. Molecular networks were exported to Cytoscape software (https://cytoscape.org/) to format the metabolic categories.

Metabolite annotation was performed in three consecutive steps. First, the ‘custom database search’ module from Mzmine was used to compare the obtained LC-MS/MS data with the IJPB chemistry and metabolomic platform homemade libraries containing 166 standards (m/z absolute tolerance of 0.005 Da and RT tolerance of 0.2 min). Second, the ESI- and ESI+ metabolomic data used for molecular network analyses were searched against the available MS2 spectral libraries (Massbank NA, GNPS Public Spectral Library, NIST14 Tandem, NIH Natural Product and MS-Dial), with absolute m/z tolerance of 0.02, 4 minimum matched peaks and minimal cosine score of 0.65. Third, not-annotated metabolites that belong to molecular network clusters containing annotated metabolites from steps 1 and 2 were assigned to the same metabolic category and searched against SIRIUS+CSI:FingerID GUI and CLI software (https://bio.informatik.uni-jena.de/software/sirius/) and MS2Query - machine learning assisted library querying of MS/MS spectra (https://github.com/iomega/ms2query) used to search for analogues.

### Statistical analysis

All statistical analyses were performed in R v.4.3.1 (R Core Team, 2023).

### Phenotypic analyses

#### STB phenotypic variation in a matrix of wheat genotype mixtures

The susceptibility of each focal genotype to STB was compared between each mixture and the pure condition. The mean ratio was calculated as xij / xii, where xij is the adjusted mean of focal genotype i grown with neighboring genotype j and xii is the adjusted mean of focal genotype i grown with the same neighboring genotype i. Adjusted means are least square means calculated from the aligned and ranked linear model using the emmeans package. Means were compared using contrast tests following an aligned and ranked linear model (Equation 1). To correct for multiple testing, a Bonferroni correction was applied for each focal genotype.

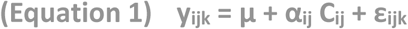

where y_ijk_ is the phenotypic observation; μ is the intercept; α_ij_ corresponds to the effect for the genotypic combination of focal i and neighbor j; C_ij_ corresponds to the genotypic combination of focal i and neighbor j; and ε_ijk_ is the error term.

#### Phenotypic analyses of Venezio

Boxplots were created to visualize STB symptoms, aerial and root phenotypes, and *Zymoseptoria tritici* gene expression from the Venezio genotype using the ggplot2, see, and stats packages. To evaluate the neighbor effect (Venezio *vs.* Soissons), Wilcoxon tests were conducted on the phenotypic observations.

### Transcriptomic analyses

#### Statistical analyses of expression data

Statistical analyses were conducted using the R script-based tool DiCoExpress Version 1 based on the Bioconductor package edgeR v 3.28.0 (Robinson *et al*., 2010; McCarthy *et al*., 2012). Two separate statistical analyses were conducted on the (i) root and (ii) leaf samples of Venezio grown in pure stands or mixed with Soissons with no treatment on the day of inoculation (0 dpi).

#### Gene filtering and normalization

For each analysis, low counts were filtered using the “filterByExpr” function where the group argument specifies the biological conditions, the min.count value is set to 15 and the min.total.count, large.n and min.prop arguments are set to their default values. Libraries were normalized with the Trimmed Mean of M-values (TMM) with the default parameter values.

#### Differential expression analysis

Transcriptomic profiles were visualized using Principal Component Analysis (PCA) (factomineR and factoextra packages), heatmaps (gplots and pheatmap packages) and scatterplots (ggplot2 package). Hierarchical clustering for heatmaps was performed using the complete clustering method and Euclidean distance, and cluster profiling was conducted using the coseq package, and visualized through boxplots (ggplot2 package).

For the two datasets, the differential analysis was based on a negative binomial generalized linear model, the logarithm of the average gene expression is an additive function of a neighbor effect (Venezio in pure or mixed with Soissons), a barrier effect (empty or filled with PVPP), their interaction, and a replicate effect. Only contrasts for the difference between the two neighbors for each type of barrier were analysed. For each contrast, a likelihood ratio test was applied, and raw p-values were adjusted with the Benjamini-Hochberg procedure to control the false discovery rate. The distribution of the resulting p-values followed the quality criterion described by Rigaill et al., 2018.

For each condition (empty or PVPP-filled barrier), log2 fold-changes between the mixed and pure conditions were calculated. Transcripts were considered significantly differentially expressed if they showed a log2 fold-change of less than −1 or greater than 1 and an adjusted p-value lower than 0.05 in the considered condition. Common differentially expressed genes between the empty and PVPP-filled barrier conditions were visualized using Venn diagrams (VennDiagram package).

To associate transcriptomic changes with phenotypic differences specific to the mixed *versus* pure stand conditions in the empty barrier but not with PVPP, transcripts were considered significantly differentially expressed if they showed a log2 fold-change of less than −1 or greater than 1 and an adjusted p-value lower than 0.05 in the empty barrier condition, while exhibiting a log2 fold-change between −1 and 1 and an adjusted p-value higher than 0.05 in the PVPP-filled barrier.

Gene Ontology (GO) term enrichments were conducted on the differentially expressed genes identified in each dataset (roots and leaves). The analysis was performed using a hypergeometric test followed by Bonferroni correction (gprofiler2 package), with the background defined as all genes expressed within each dataset. Results were visualized as a network using EnrichmentMap in Cytoscape.

### Metabolomic analyses

Metabolomic profiles were visualized using Principal Component Analysis (PCA) (factomineR and factoextra packages), heatmaps (gplots and pheatmap packages), and scatterplots (ggplot2 package). Hierarchical clustering for heatmaps was conducted using the complete clustering method with Euclidean distance, and cluster profiling was performed using the co-seq package, visualized through boxplots (ggplot2 package).

#### Leaf & root metabolomic analyses

For metabolite analysis of the two datasets (roots and leaves) at 0 dpi, log2 fold-changes between the mixed and pure conditions and z-scores (Equation 2) were calculated for each condition (empty barrier and barrier with PVPP).

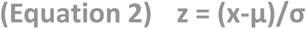

where z is the vector of z-scores; x is the log2 fold-change of metabolite accumulation between mixture and pure condition; μ is the mean of log2 fold-changes; and σ is the standard error of the log2 fold-changes.

Metabolites were considered significantly differentially accumulated if they showed a log2 fold-change of less than −1 or greater than 1 and a z-score below −2 or above 2 in the considered condition. Common differentially accumulated metabolites between the empty and PVPP-filled barrier conditions were visualized using Venn diagrams (VennDiagram package).

To link metabolite changes with phenotypic differences between the mixed and pure conditions in the empty barrier but not with PVPP, metabolites were considered significantly differentially accumulated if they exhibited a log2 fold-change less than −1 or greater than 1 and a z-score below −2 or above 2 in the empty barrier, while displaying a log2 fold-change between −1 and 1 and a z-score between −2 and 2 in the PVPP-filled barrier.

#### Soil metabolomic analyses

PLS-DA coordinates from soil samples were used to test interactions between neighbor (Venezio *vs.* Soissons) and barrier condition (empty barrier and barrier with PVPP) using a non-parametric ANOVA (Equation 3):

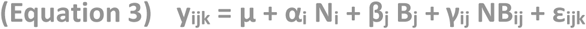

where y_ijk_ is the PLS-DA coordinate; μ is the intercept; α_i_, β_j_ and γ_ij_ correspond to effects for neighbor i, barrier condition j and interaction between neighbor i and barrier j, respectively; N_i_, B_j_ and NB_ij_ correspond to neighbor i, barrier condition j and interaction between neighbor i and barrier condition j, respectively; and ε_ijk_ is the error term.

For soil metabolite analysis at 0 dpi, log2 fold-changes between the mixed and pure conditions and z-scores (Equation 2) were calculated for each barrier condition (empty barrier and barrier with PVPP). Metabolites were considered significantly differentially accumulated if they exhibited a log2 fold-change below −1 or above 1 and a z-score less than −2 or greater than 2 in the empty barrier, while displaying a log2 fold-change between −1 and 1 and a z-score between −2 and 2 in the PVPP-filled barrier.

## Author contributions

LM conceived and designed the experiments. LM, AC, and SM performed aerial phenotyping. LM, AC, and JS conducted root phenotyping. LM and AC performed the multi-omic experiments and RNA extractions. ALA, CPLR and MLM carried out RNA sequencing and preliminary data analyses. JCT, FP, CR, AR, and PP generated the metabolomic data. LM conducted all computational analyses and prepared the figures and tables. LM led the writing of the manuscript. LVM, JBM, and EB supervised the work. LM, LVM, JBM, and EB reviewed the manuscript. All authors read and approved the final version.

## Acknowledgments

We kindly thank Sandrine Roques for providing the rhizoboxes that were used for root phenotyping.

This work was supported by the French National Research Agency under the Investments for the Future Program [grants MOBIDIV project ANR-20-PCPA-0006 and TRADEOFF project ANR-19-CE20-0005]. It also benefited from the support of the Bordeaux Metabolome Facility and the IJPB Plant Observatory platform PO-Chem. The Bordeaux Metabolome Facility was funded by the MetaboHUB (ANR-11-INBS-0010) and PHENOME (ANR-11-INBS-0012) projects. The IJPB additionally benefits from the Saclay Plant Sciences-SPS program (ANR-17-EUR-0007).

## Data availability statement

All datasets and code used for data processing, analysis, and figure reproduction are available in the Zenodo repository (https://doi.org/10.5281/zenodo.17916677). Untargeted metabolomic raw data have been deposited in the MassIVE repository (https://massive.ucsd.edu/ProteoSAFe/static/massive.jsp) under the following identifiers: (MSV000100170) for soil, (MSV000100171) for leaf and (MSV000100169) for root datasets. Transcriptomic RNA-Seq raw FASTQ files are accessible through the NCBI Sequence Read Archive (https://www.ncbi.nlm.nih.gov/sra/) under BioProject accession (PRJNA1379485).

## Supplemental information

**Figure S1.**
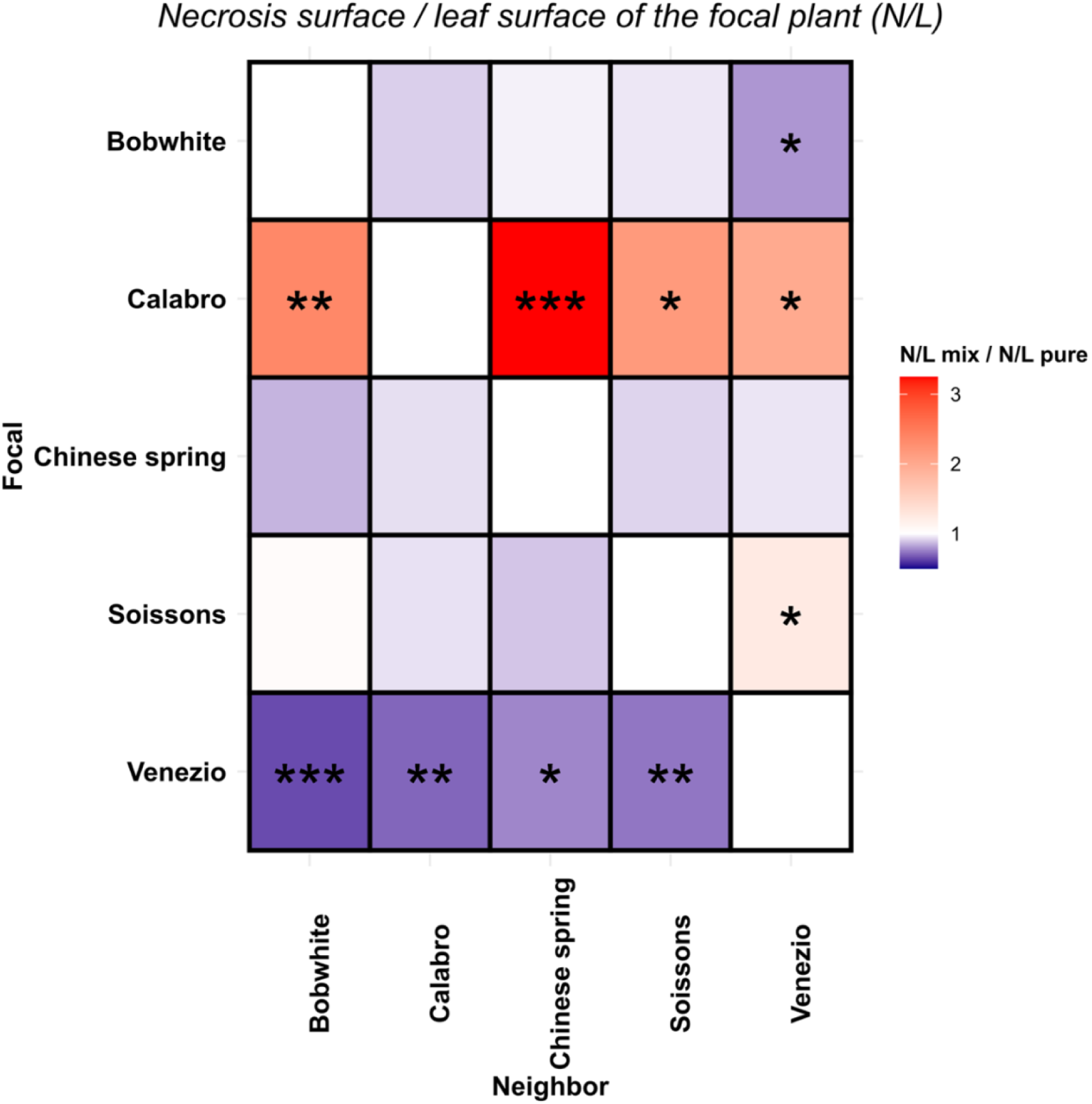
Modulation of focal average susceptibility to Septoria tritici blotch (STB) by neighboring plants. The ratio of focal necrosis surface in each mixture compared to pure condition was calculated and depicted in the Figure; thus diagonal values are equal to 1. Cases where the focal susceptibility in mixture is higher than pure condition are indicated in red, while those with lower susceptibility are shown in blue. Statistical comparisons between focal susceptibility in each mixture and pure condition were conducted using non-parametric contrast tests. Significance levels are denoted as follows:.: p < 0.1, *: p < 0.05, **: p < 0.01 and ***: p < 0.001.

**Figure S2.**
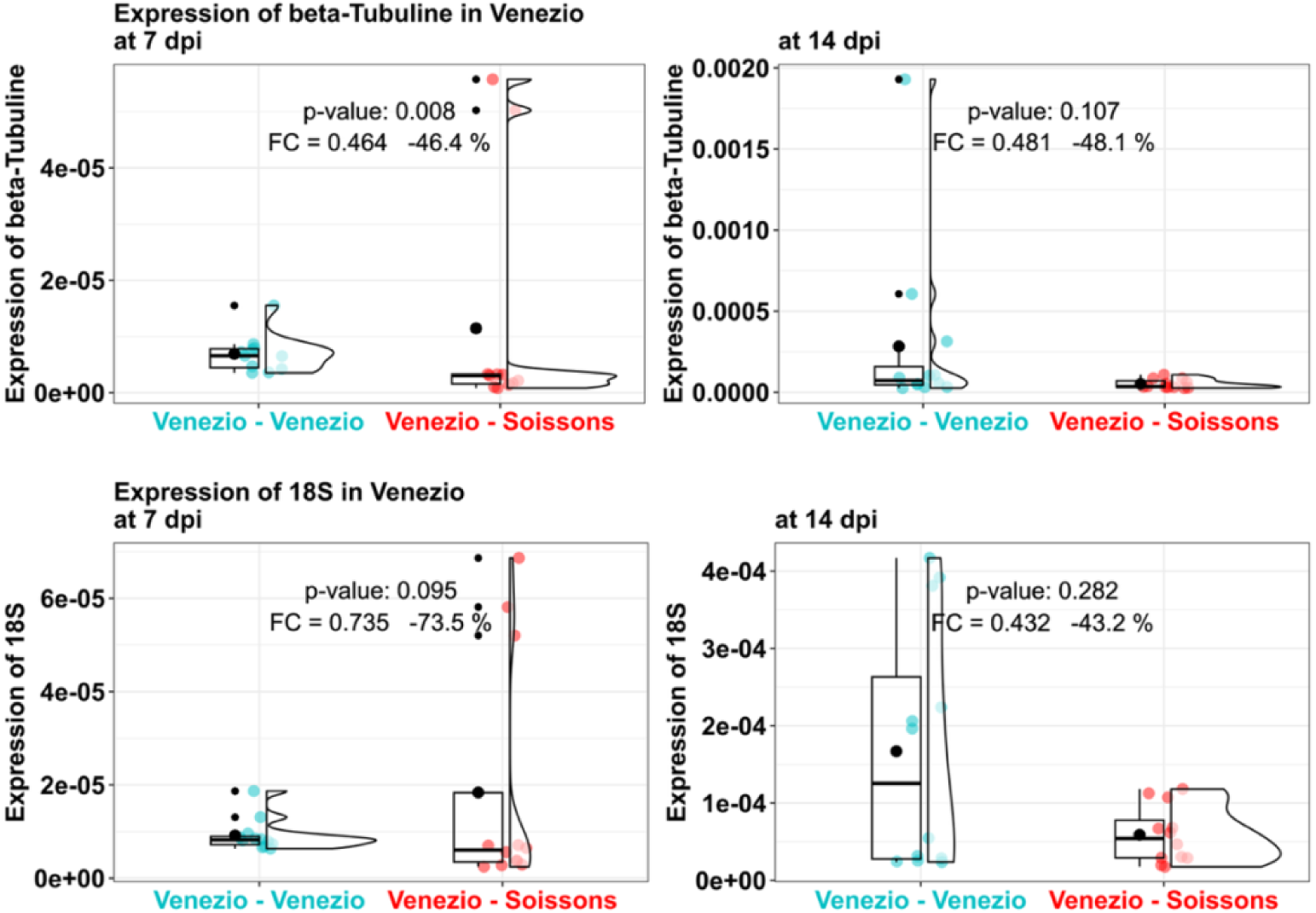
*Zymoseptoria tritici* gene expression quantification in focal leaves grown in pure stands *vs.* in a bread wheat cultivar mixture. Comparison of gene expression levels in inoculated Venezio leaves grown in pure stands *versus* mixed with Soissons at 7 days (left panel) and 14 days (right panel) post-inoculation. Boxplots show the expression levels of the pathogen housekeeping genes (18S and β-tubuline). Statistical analyses between pure and mixed conditions were conducted using non-parametric ANOVA.

**Figure S3.**
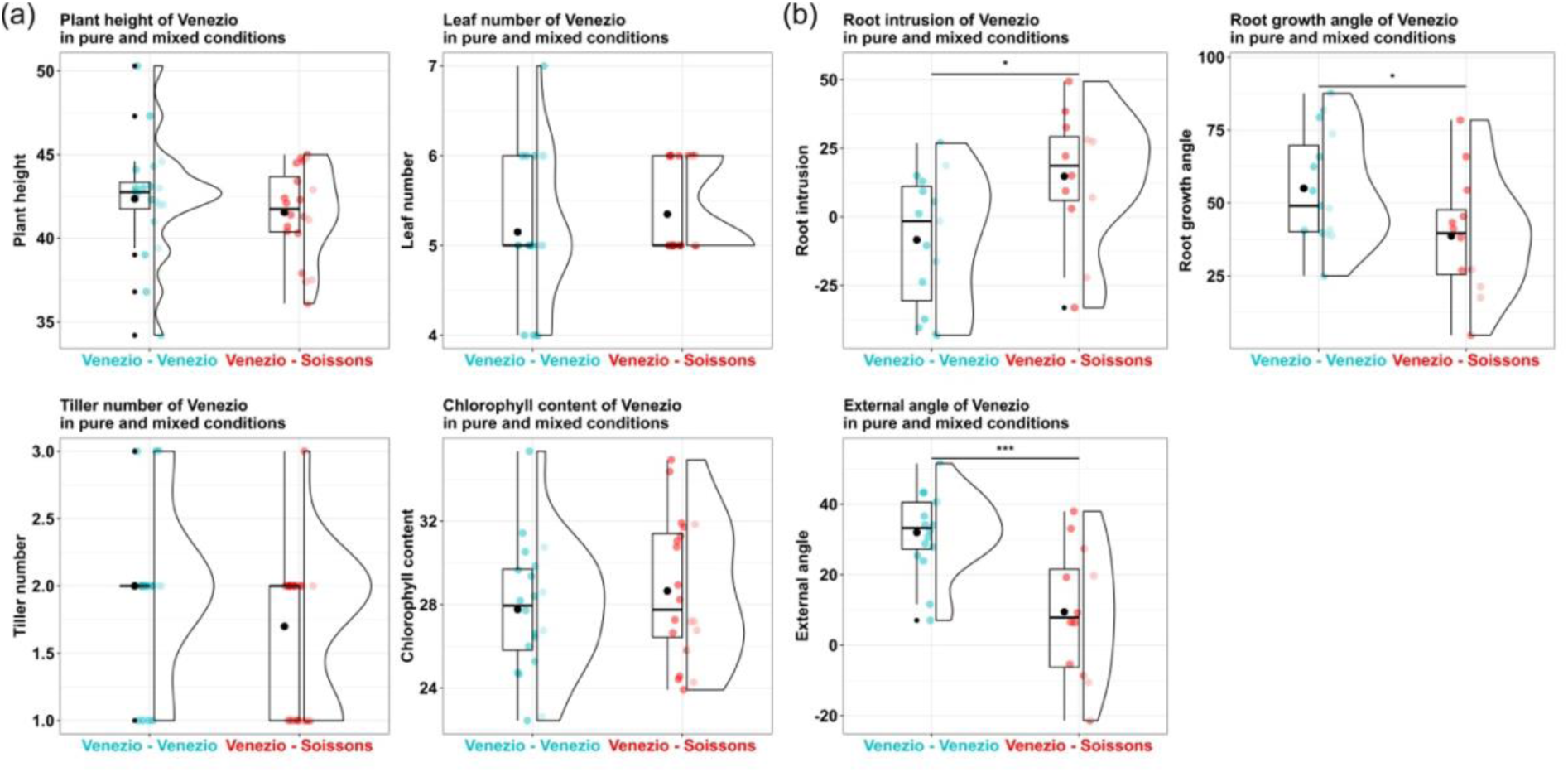
Effect of neighbor genotype on aerial and root traits in a bread wheat cultivar mixture. (a) Aerial and (b) root phenotyping of the Venezio genotype was assessed 38 and 21 days after germination, respectively. Plants were grown either in pure stands or in mixtures with Soissons. Statistical analyses between pure and mixed conditions were conducted using non-parametric contrast tests.

**Figure S4.**
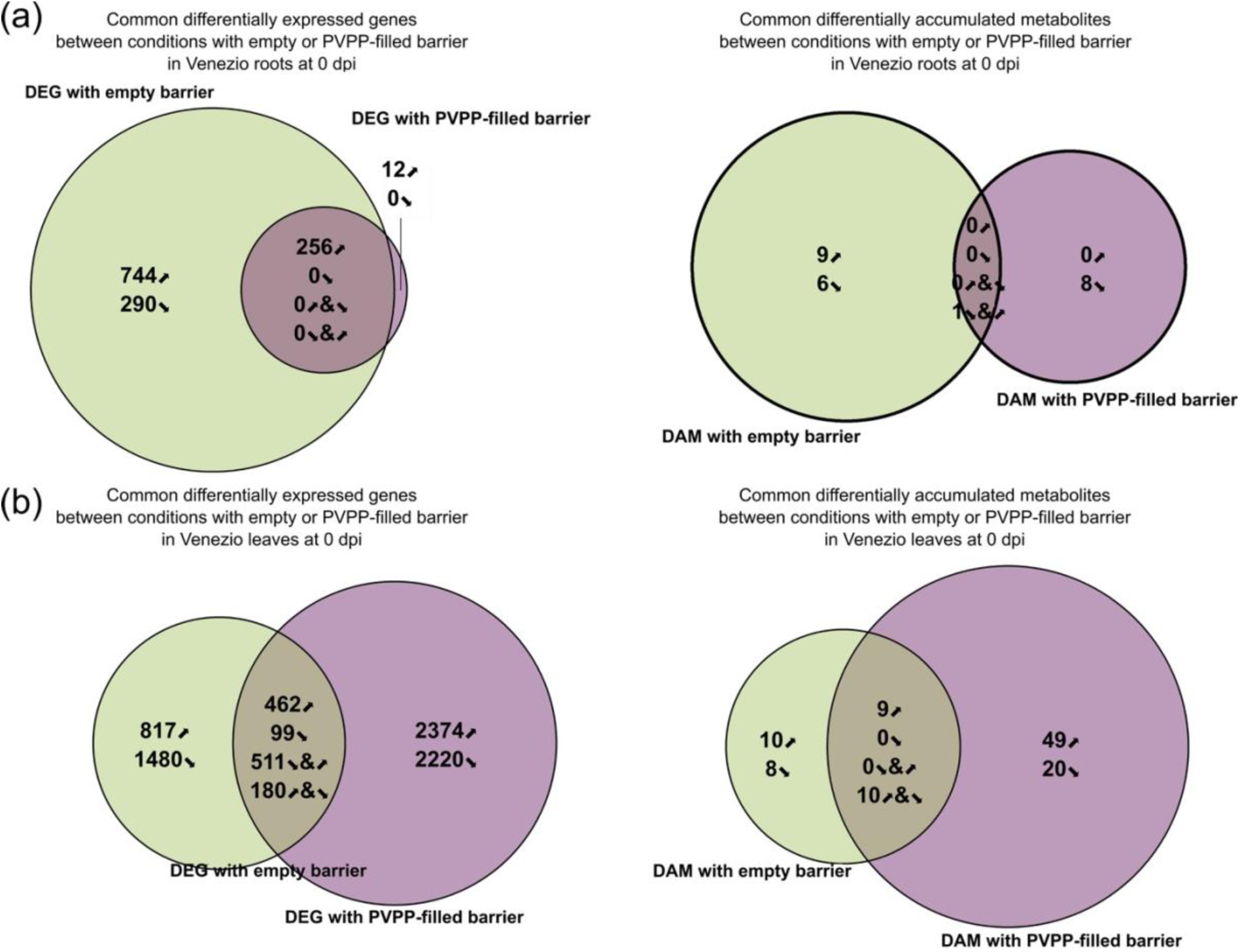
Differentially expressed genes (DEG) and differentially accumulated metabolites (DAM) between pure and mixed conditions under empty and PVPP-filled barrier conditions. Shown are all DEG (left panel) and DAM (right panel) between pure and mixed conditions with Soissons for Venezio (a) roots and (b) leaves at 21 days after germination (the day of inoculation), when grown with an empty barrier (green) or with a barrier containing polyvinylpolypyrrolidone (PVPP; purple). Overlapping regions of the circles indicate DEGs or DAMs shared between barrier conditions. Genes were considered significantly differentially expressed if they showed a log2 fold-change less than −1 or greater than 1 and an adjusted p-value lower than 0.05 in the considered condition. Metabolites were classified as significantly differentially accumulated if they showed a log2 fold-change less than −1 or greater than 1 and a z-score below −2 or above 2 in the considered condition.

**Figure S5.**
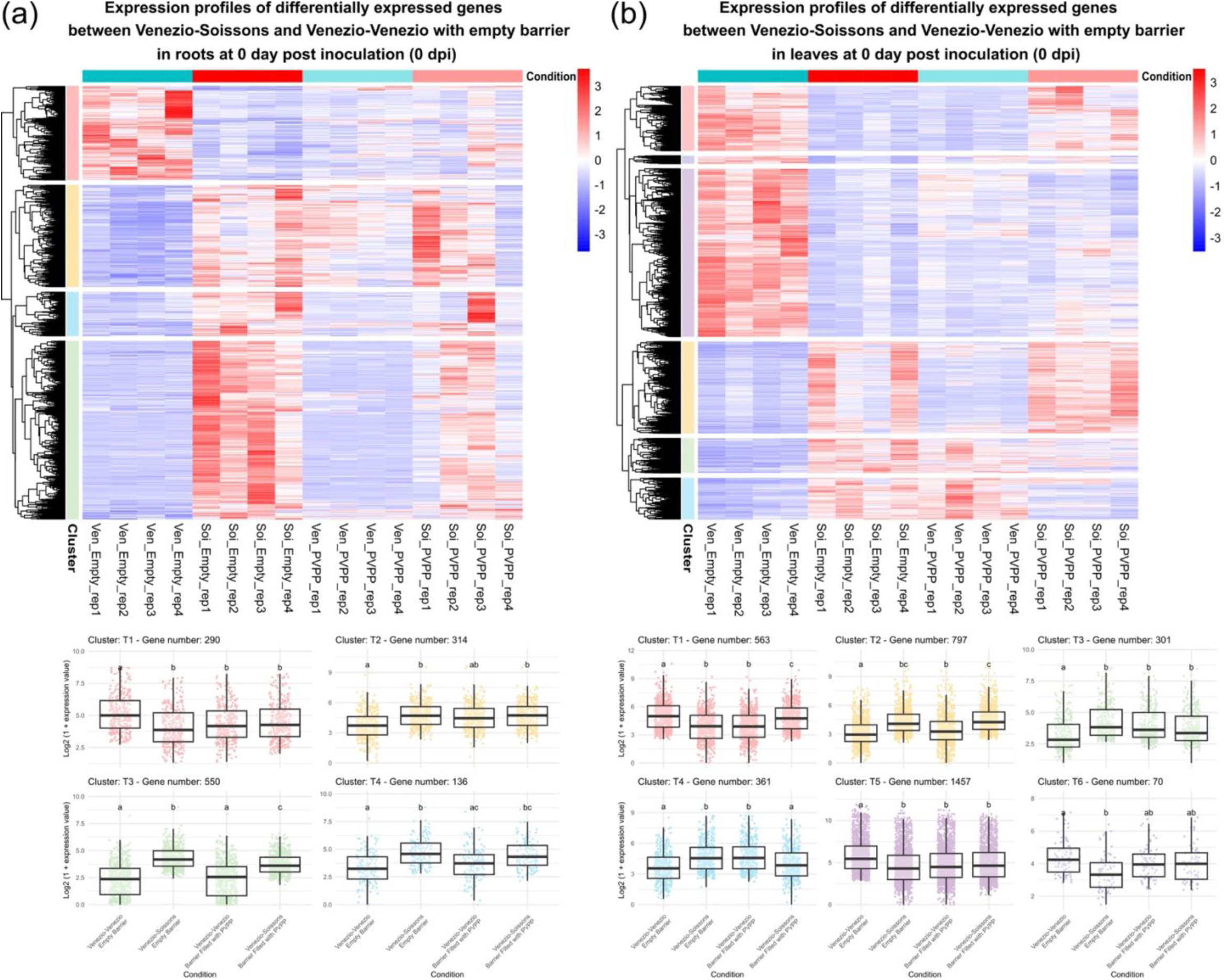
Expression profiles of differentially expressed genes in pure and mixed Venezio when grown with an empty barrier. Expression profiles and associated clusters are shown or differentially expressed genes in Venezio (a) roots and (b) leaves at 21 days after germination (the day of inoculation), between pure and mixed conditions with Soissons, when grown with an empty barrier. Genes were considered significantly differentially expressed if they showed a log2 fold-change of less than −1 or greater than 1 and an adjusted p-value ower than 0.05 in the empty barrier condition.

**Figure S6.**
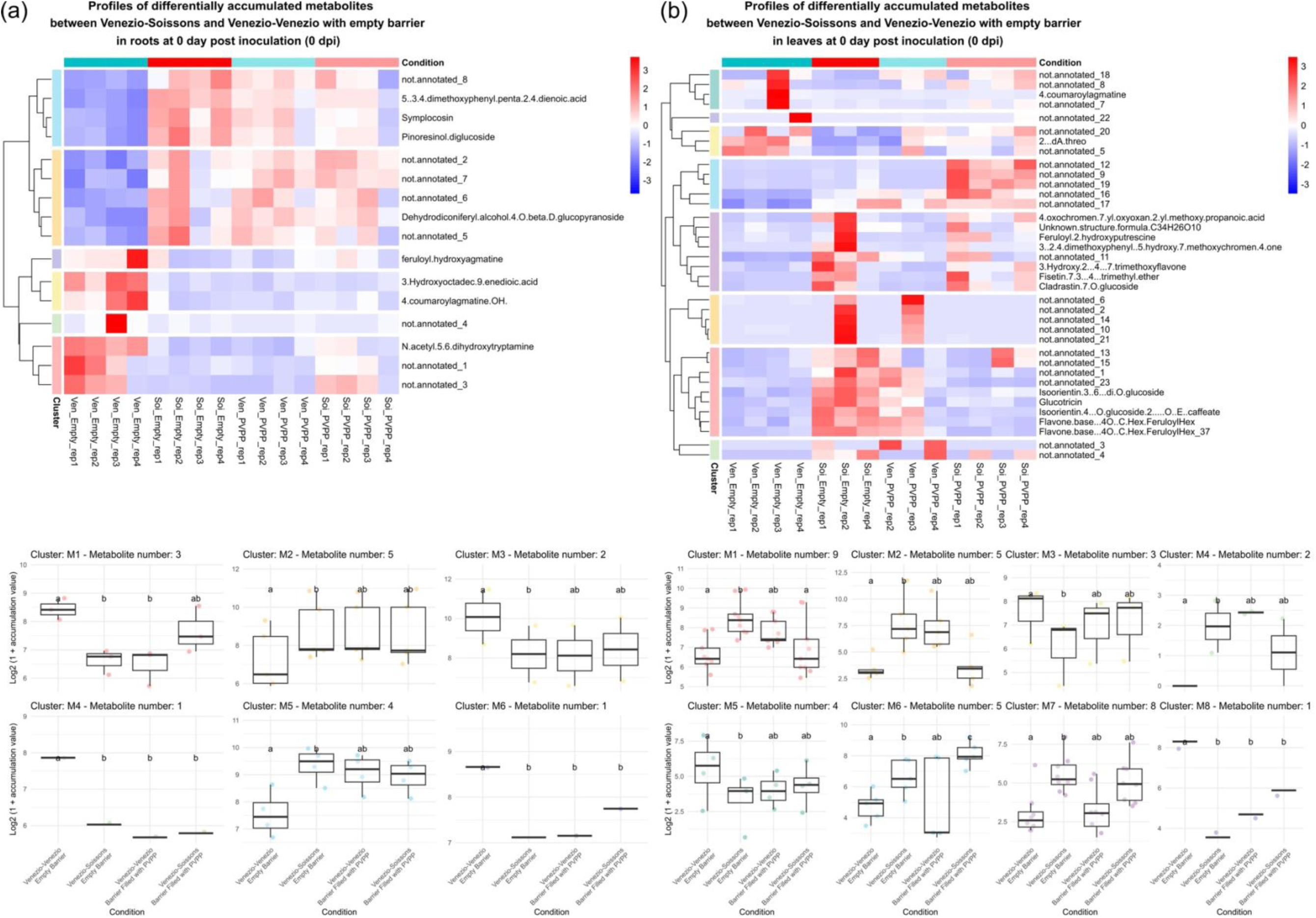
Accumulation profiles of differentially accumulated metabolites in pure and mixed Venezio when grown with an empty barrier. Accumulation profiles and associated clusters are shown for differentially accumulated metabolites in Venezio (a) roots and (b) leaves at 21 days after germination (the day of inoculation), between pure and mixed conditions with Soissons, when grown with an empty barrier. Metabolites were considered significantly differentially accumulated if they exhibited a log2 fold-change less than-1 or greater than 1 and a z-score below −2 or above 2 in the empty barrier condition.

**Figure S7.**
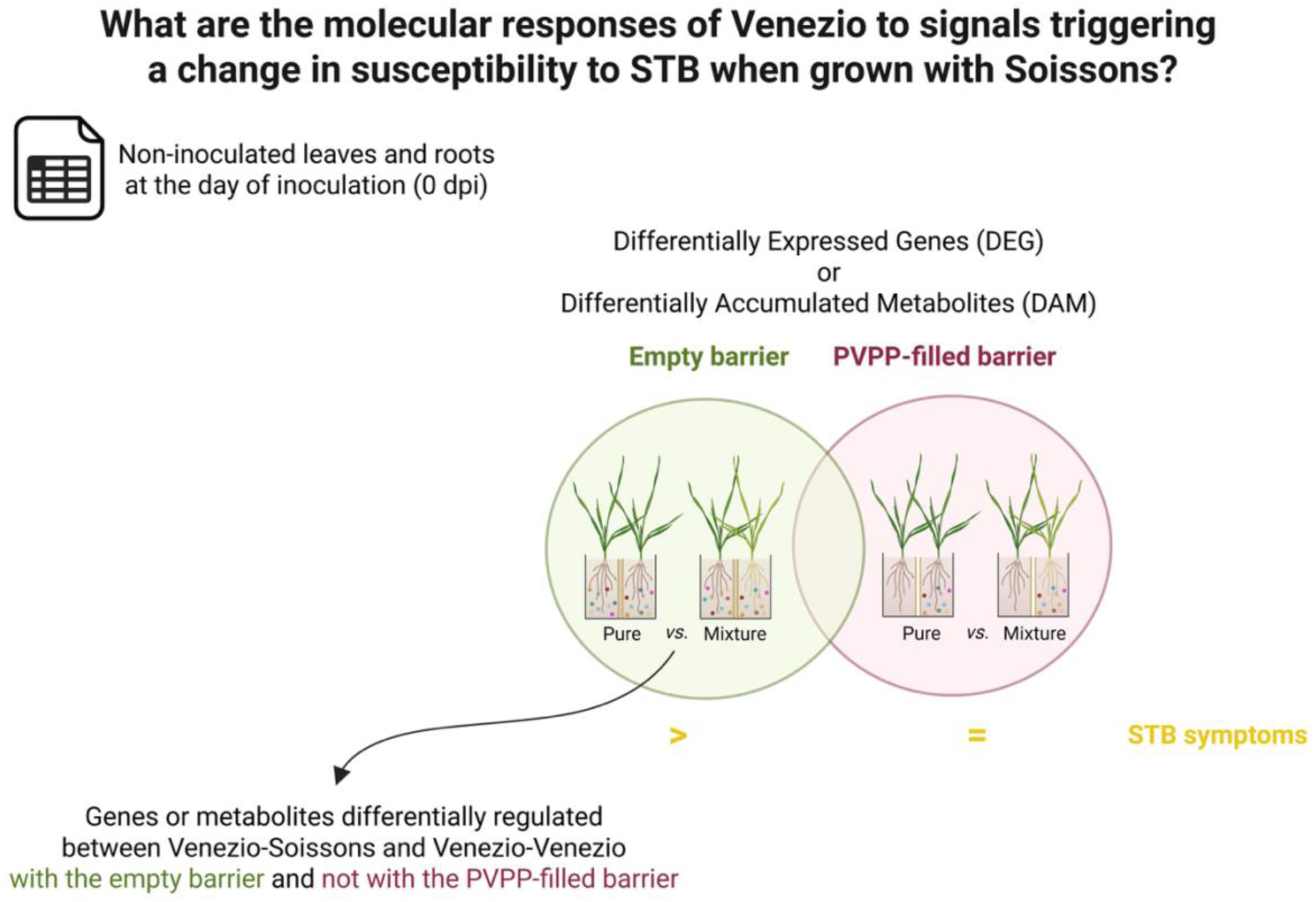
Overview of the experimental design and analytical workflow for studying the modulation of *Zymoseptoria tritici* susceptibility in a bread wheat cultivar mixture. Using the Venezio–Soissons cultivar mixture as a model system, we investigated the molecular responses of bread wheat (cv. Venezio) to signals that reduce susceptibility to Septoria tritici blotch (STB) when grown in mixture. Transcriptomic and metabolomic analyses were performed on leaves and roots of Venezio plants harvested at 21 days after germination (day of inoculation). Plants were grown either in pure stands or in mixture with Soissons, separated by one of two types of physical barriers: an empty barrier, or a barrier filled with polyvinylpolypyrrolidone (PVPP), a compound that adsorbs phenolic molecules. Genes and metabolites were selected based on differential regulation between pure and mixed conditions only in the presence of the empty barrier, but not with the PVPP-filled barrier. This approach enabled the identification of molecular features potentially associated with reduced STB susceptibility in mixtures, an effect observed with the empty barrier but abolished when PVPP was present.

**Figure S8.**
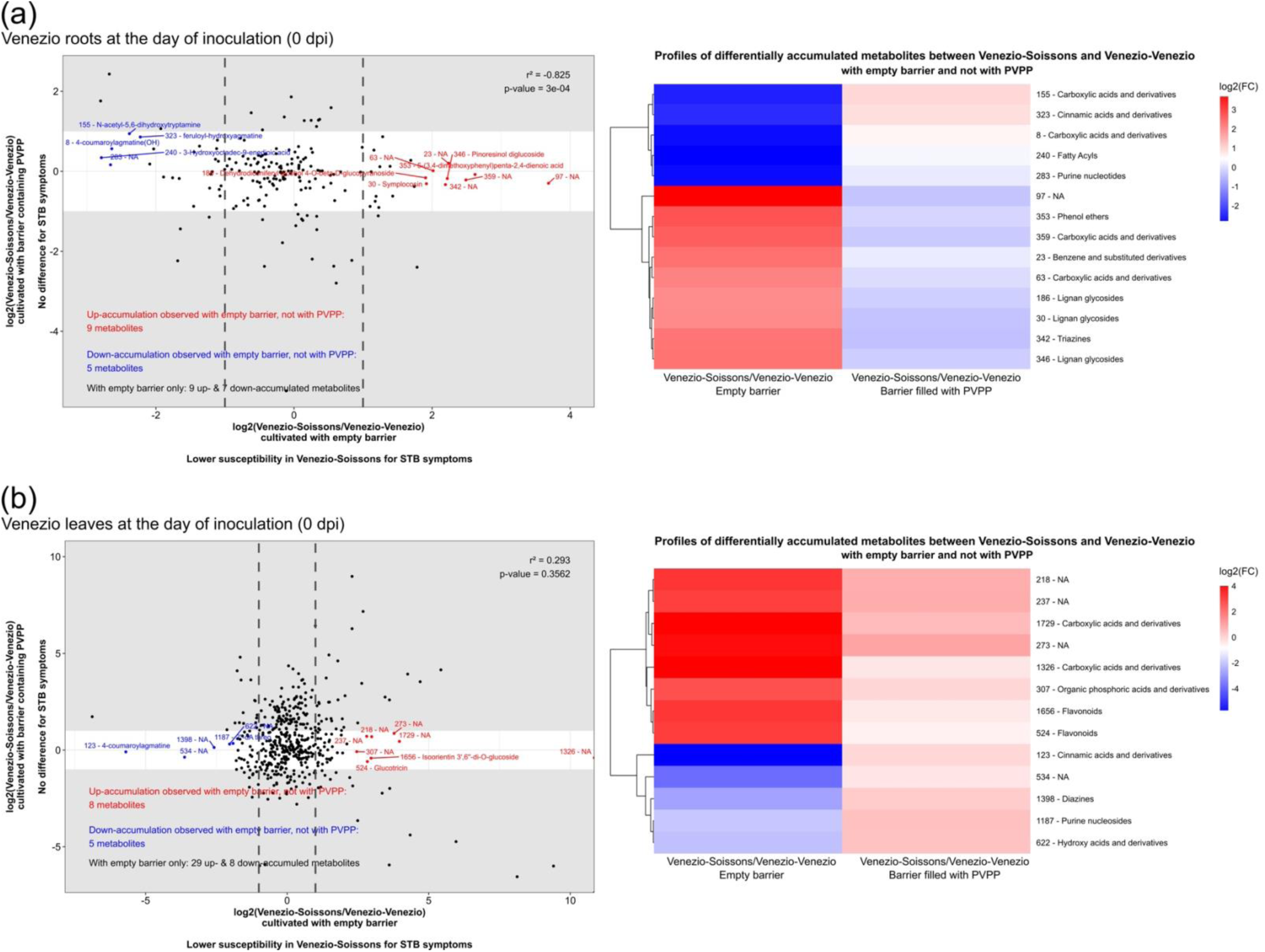
Accumulation profiles of differentially accumulated metabolites between pure or mixed Venezio when grown with an empty barrier, and not with a PVPP-filled barrier. Accumulation profiles and associated metabolite classes are shown for differentially accumulated metabolites in Venezio (a) roots and (b) leaves at 21 days after germination (day of inoculation), between pure and mixed conditions with Soissons, when grown with an empty barrier, and not with a barrier containing polyvinylpolypyrrolidone (PVPP). Metabolites were considered significantly differentially accumulated if they exhibited a log2 fold-change less than −1 or greater than 1 and a z-score below −2 or above 2 in the empty barrier condition, while displaying a log2 fold-change between −1 and 1 and a z-score between −2 and 2 with the PVPP-filled barrier.

**Figure S9.**
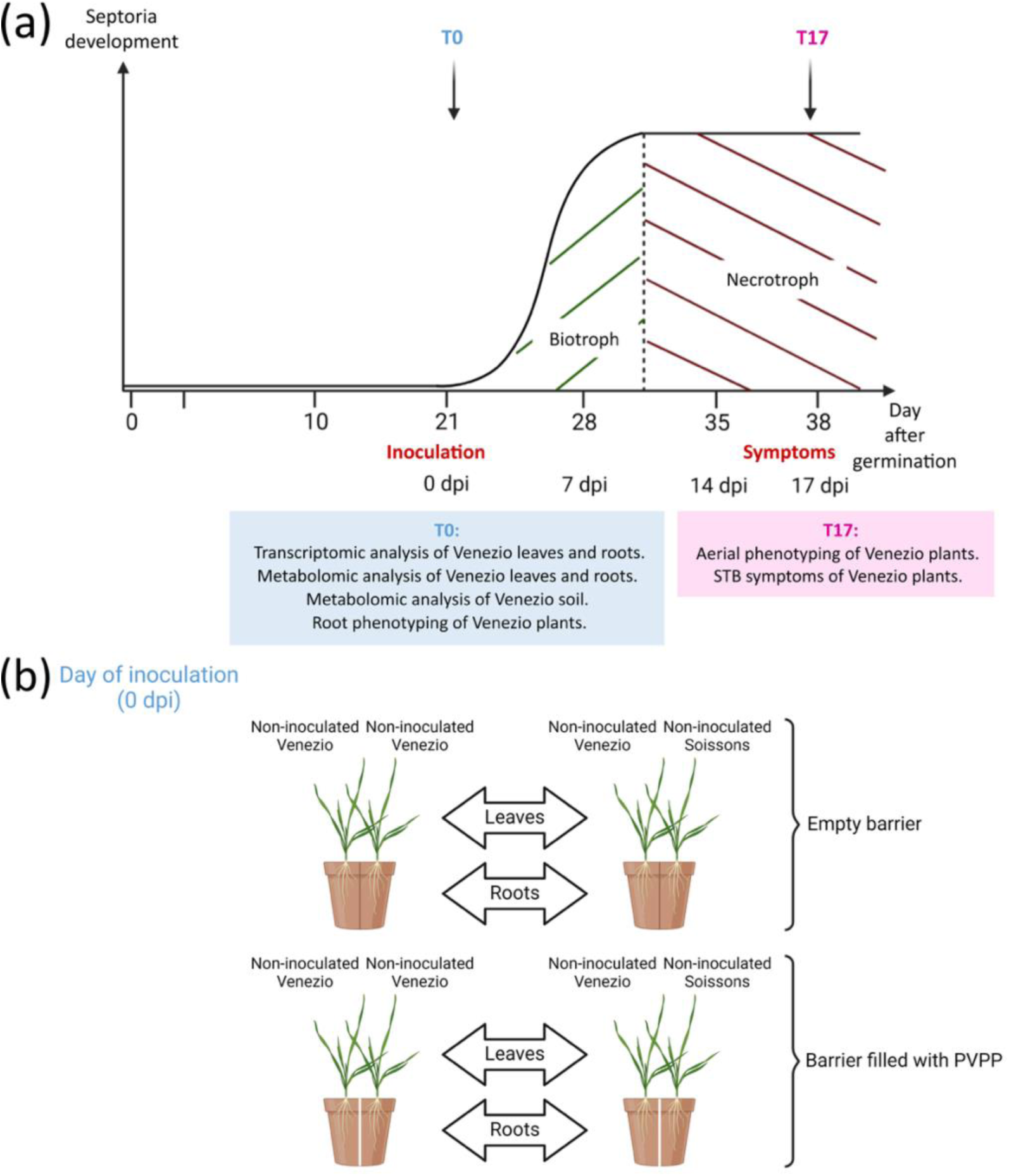
Experimental design for studying the modulation of *Zymoseptoria tritici* susceptibility in a bread wheat cultivar mixture. (a) 21 days after germination (day of inoculation): Transcriptomic and metabolomic analyses were conducted on both leaves and roots of Venezio. Metabolomic analysis of soil associated with Venezio was performed, comparing samples from pure stands and mixed with Soissons when cultivated with an empty barrier or a barrier containing polyvinylpolypyrrolidone (PVPP) between genotypes. Root phenotyping was carried out to compare the root architecture of Venezio between pure and mixed conditions. 17 days post-inoculation: Aerial phenotyping and assessment of disease symptoms were conducted on Venezio to compare the level of *Zymoseptoria tritici* susceptibility between plants grown in pure stands *versus* mixed with Soissons when cultivated under control conditions, with an empty barrier or a barrier containing PVPP. (b) Transcriptomic and metabolomic analyses were conducted on non-inoculated leaves and roots of Venezio at 21 days after germination (day of inoculation) to compare responses between pure stands and mixtures with Soissons, under conditions with either an empty barrier or a barrier containing PVPP.

**Table S1** Detailed list of detected metabolites in soil. For each metabolite, the table provides the name, superclass, class and subclass, along with the pathways it is involved in, its KEGG ID, and RefMetaDB ID. Metabolites identified within the interaction clusters are indicated.

**Table S2** Datasets used to perform the multi-omic analyses. Number of genes and metabolites available in each dataset.

**Table S3** Number of genes and metabolites differentially regulated in each dataset. Number of up-regulated (↗) and down-regulated (↘) genes or metabolites in Venezio grown in mixture compared to pure conditions under three settings: with an empty barrier, with a barrier containing polyvinylpolypyrrolidone (PVPP) or with an empty barrier but not with a PVPP-filled barrier.

**Table S4** Detailed list of differentially expressed genes between pure or mixed Venezio roots when grown with an empty barrier, but not with a PVPP-filled barrier, at 21 days after germination (day of inoculation). (a) Up- and (b) down-expressed genes in the mixture with Soissons compared to the pure condition in the empty barrier condition are listed. Genes were considered significantly differentially expressed if they showed a log2 fold-change of less than −1 or greater than 1 and an adjusted p-value lower than 0.05 in the empty barrier condition, while exhibiting a log2 fold-change between −1 and 1 and an adjusted p-value higher than 0.05 with the PVPP-filled barrier.

**Table S5** Detailed list of differentially expressed genes between pure or mixed Venezio leaves when grown with an empty barrier, but not with a PVPP-filled barrier, at 21 days after germination (day of inoculation). (a) Up- and (b) down-expressed genes in the mixture with Soissons compared to the pure condition in the empty barrier condition are listed. Genes were considered significantly differentially expressed if they showed a log2 fold-change of less than −1 or greater than 1 and an adjusted p-value lower than 0.05 in the empty barrier condition, while exhibiting a log2 fold-change between −1 and 1 and an adjusted p-value higher than 0.05 with the PVPP-filled barrier.

**Table S6** Detailed list of differentially accumulated metabolites between pure or mixed Venezio roots when grown with an empty barrier, but not with a PVPP-filled barrier, at 21 days after germination (day of inoculation). (a) Up- and (b) down-accumulated metabolites in the mixture with Soissons compared to the pure condition in the empty barrier condition are listed. Metabolites were considered significantly differentially accumulated if they exhibited a log2 fold-change less than −1 or greater than 1 and a z-score below −2 or above 2 in the empty barrier condition, while displaying a log2 fold-change between −1 and 1 and a z-score between −2 and 2 with the PVPP-filled barrier.

**Table S7** Detailed list of differentially accumulated metabolites between pure or mixed Venezio leaves when grown with an empty barrier, but not with a PVPP-filled barrier, at 21 days after germination (day of inoculation). (a) Up- and (b) down-accumulated metabolites in the mixture with Soissons compared to the pure condition in the empty barrier condition are listed. Metabolites were considered significantly differentially accumulated if they exhibited a log2 fold-change less than −1 or greater than 1 and a z-score below −2 or above 2 in the empty barrier condition, while displaying a log2 fold-change between −1 and 1 and a z-score between −2 and 2 with the PVPP-filled barrier.

**Table S8** Detailed list of genes used for analyzing transcriptional changes in *Zymoseptoria tritici*. For each gene, the table provides the gene name and primer sequences.

